# Proliferative behavior of hematopoietic stem cells revisited: No evidence for mitotic memory

**DOI:** 10.1101/745729

**Authors:** Mina N. F. Morcos, Thomas Zerjatke, Ingmar Glauche, Clara M. Munz, Yan Ge, Andreas Petzold, Susanne Reinhardt, Andreas Dahl, Natasha Anstee, Ruzhica Bogeska, Michael Milsom, Petter Säwén, Haixia Wan, David Bryder, Axel Roers, Alexander Gerbaulet

## Abstract

The proliferative activity of adult hematopoietic stem cells (HSCs) is controversially discussed. Inducible fluorescent histone 2B fusion protein (H2B-FP) transgenic mice are important tools for tracking the mitotic history of murine HSCs in label dilution experiments. A recent study proposed that the most primitive HSCs divide only four times, to then enter permanent quiescence. We observed that background fluorescence due to leaky H2B-FP expression, occurring in all H2B-FP transgenes independent of label induction, accumulated with age in primitive HSCs with high repopulation potential. We argue that this background had been misinterpreted as retention of induced label and permanent quiescence. We found cell division-independent half-lives of H2B-FPs to be short, which had led to overestimation of HSC divisional activity. Our data do not support HSC mitotic memory and entry into permanent quiescence after few divisions, but show that primitive HSCs of adult mice continue to cycle rarely.

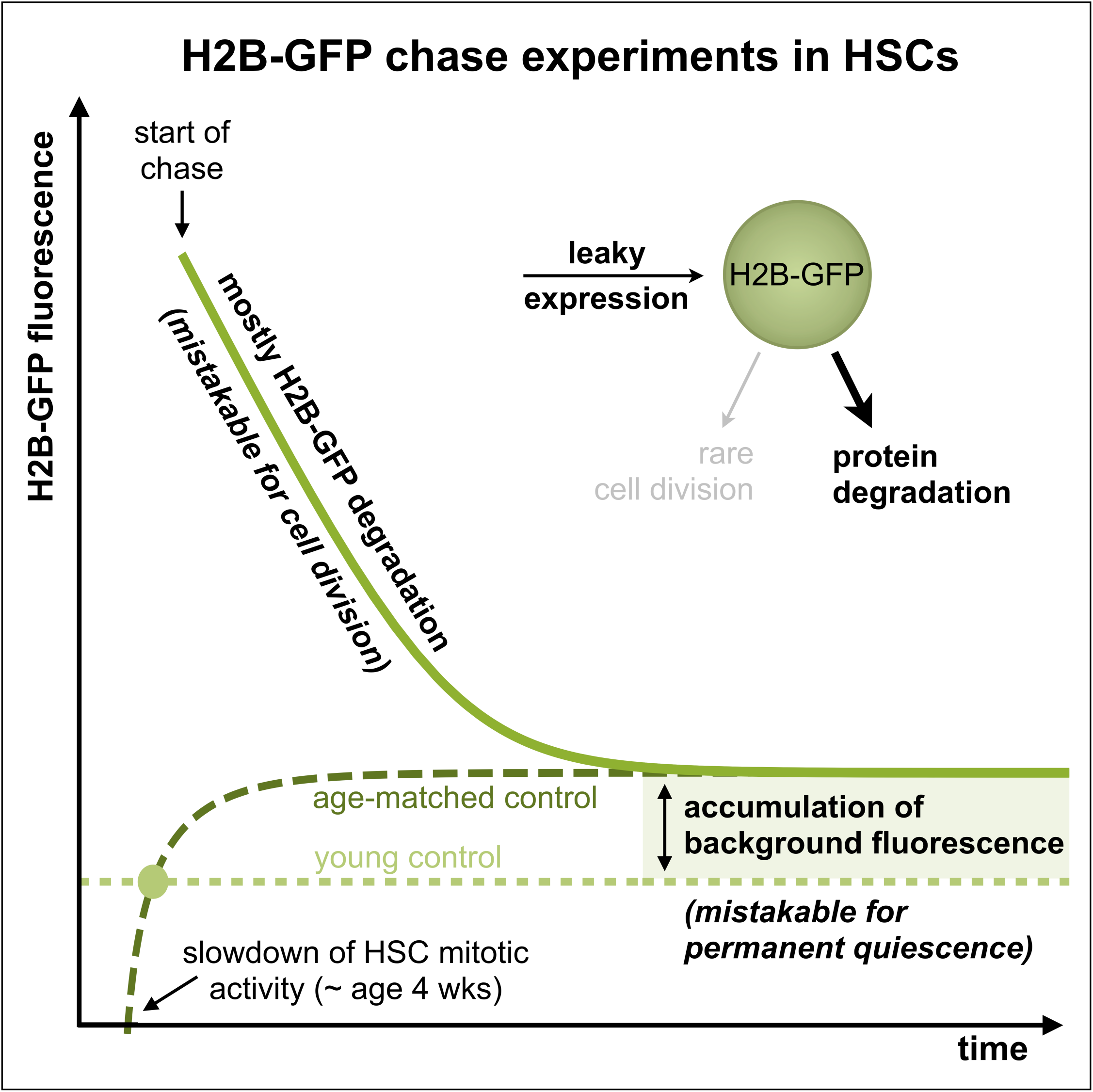

## Introduction

The multi-potent and self-renewing hematopoietic stem cells (HSCs) reside at the top of the hematopoietic hierarchy and can give rise to all blood lineages throughout the life of an individual (Eaves, 2015). Numerous studies have demonstrated heterogeneous cell cycle activity within the HSC population (Foudi et al., 2008; Glauche et al., 2009; Morcos et al., 2017; Passegué et al., 2005; Qiu et al., 2014; Säwén et al., 2016; Takizawa et al., 2011; Wilson et al., 2008). A concept of deeply quiescent (also termed ‘dormant’) and actively cycling HSCs has been inferred from experiments investigating mitotic history by means of labeled nucleotide analogs (e.g. BrdU) (Kiel et al., 2007; Wilson et al., 2008) or histone 2B (H2B) fused to either enhanced green fluorescent protein (H2B-GFP) or mCherry (hereafter referred to as H2B-RFP) (Foudi et al., 2008; Qiu et al., 2014; Säwén et al., 2016; Wilson et al., 2008). These labels can be incorporated into chromatin and get homogenously distributed to daughter cells during mitosis (Tumbar et al., 2004), which allows tracking several cell divisions as well as identifying quiescent label retaining cells. In case of the H2B-fusion proteins (H2B-FPs), labeling of HSCs is achieved with inducible tetracycline (Tet) controlled (either ‘Tet-on’ or ‘Tet-off’) genetic systems (Gossen and Bujard, 2002). Importantly, these mouse models exhibit considerable background fluorescence even in the repressed state due to leaky expression from the Tet-operon (Challen and Goodell, 2008; Foudi et al., 2008; Qiu et al., 2014; Säwén et al., 2016; Tumbar et al., 2004; Wilson et al., 2008).

A recent study (Bernitz et al., 2016) reported HSCs retaining H2B-GFP label for up to 22 months of chase. Based on this finding, the authors proposed a model in which the most primitive HSCs that contain all long-term repopulating activity enter complete and permanent quiescence after initially undergoing four symmetric self-renewal divisions. However, neither division-independent loss by protein degradation (Waghmare et al., 2008), which even in case of slow degradation would preclude long chase periods, nor the continuous leaky background expression (Challen and Goodell, 2008) of H2B-FPs were considered in this model.

In the present study, we re-evaluated H2B-FP retention. We show that in three different H2B-FP transgenic mouse strains, the most quiescent HSC sub-population inevitably accumulates high background fluorescence in an age-dependent manner. Accordingly, long-term serial repopulation activity was enriched among cells with high background. We estimated division-independent degradation of H2B-GFP and H2B-RFP to proceed in HSCs with half-lives of approximately 4-6 and 2 weeks, respectively, and show that neglect of H2B-FP decay leads to overestimation of HSC mitotic activity. Two sequential rounds of H2B-RFP induction and subsequent dilution revealed that HSCs do not abruptly halt mitotic activity upon ageing, arguing against permanent quiescence of HSCs after four divisions as previously hypothesized (Bernitz et al., 2016). Mathematical modeling of H2B-FP systems revealed that background label accumulates over time in the most primitive and quiescent HSCs, providing an explanation for observing seemingly label-retaining HSCs after extended chase periods. We argue that such cells were previously misinterpreted as permanently quiescent HSCs.

## Results

### Primitive HSPCs exhibit higher levels of leaky H2B-FP background fluorescence

Upon flow cytometric analysis of HSPC populations (Figure S1A) isolated from bone marrow (BM) of deliberately un-induced *R26*^rtTA^*/Col1A1*^H2B-RFP^ (Egli et al., 2007)*, R26*^rtTA^*/Col1A1*^H2B-GFP^ (Foudi et al., 2008) or single transgenic tetO-H2B-GFP47Efu (Bernitz et al., 2016; Tumbar et al., 2004) mice, we observed a gradual increase of median and maximum (as judged by the 99^th^ percentile) background H2B-FP fluorescence from fast cycling hematopoietic progenitor cells (HPCs) to quiescent HSCs (Figure 1A-C). Interestingly, within the immuno-phenotypic HSC population (LSK CD48^−/lo^CD150^+^), we found particularly bright H2B-FP background fluorescence in the predominantly quiescent CD201^hi^Sca-1^hi^CD34^−/lo^ (ES34) HSC subpopulation. Among HSCs with high background fluorescence from all three different repressed H2B-FP mouse strains under investigation, we observed significantly higher expression of Sca-1, CD201 and CD150 as well as concordant down-regulation of CD34, CD48 and CD117 (Figure S1C-E) compared to cells without H2B-FP background fluorescence. This expression pattern of surface markers has been previously correlated to increased quiescence and repopulation activity of HSCs (Beerman et al., 2010; Grinenko et al., 2014; Kent et al., 2009; Kiel et al., 2005; Morcos et al., 2017; Osawa et al., 1996; Sato et al., 1999; Säwén et al., 2016; Shin et al., 2014; Wilson et al., 2015). To further investigate the cell cycle status of HSPCs in relation to H2B-GFP background fluorescence, we analyzed the BM of un-induced *Ki67*^RFP/wt^*/R26*^rtTA/wt^*/Col1A1*^H2B-GFP/wt^ mice, in which Ki67-RFP expression reports cycling cells (Basak et al., 2014; Morcos et al., 2017). We observed a significantly lower fraction of cycling Ki67-RFP^+^ cells among HSCs and MPPs with high H2B-GFP background (Figure 1D). In addition, we performed transcriptome analysis of 200 single HSCs, which were index-sorted from un-induced *R26*^rtTA/rtTA^*/Col1A1*^H2B-GFP/H2B-GFP^ animals and found a quiescence-related gene signature upregulated in HSCs with high H2B-GFP background fluorescence (Figure S1F-J).

**Figure 1.**
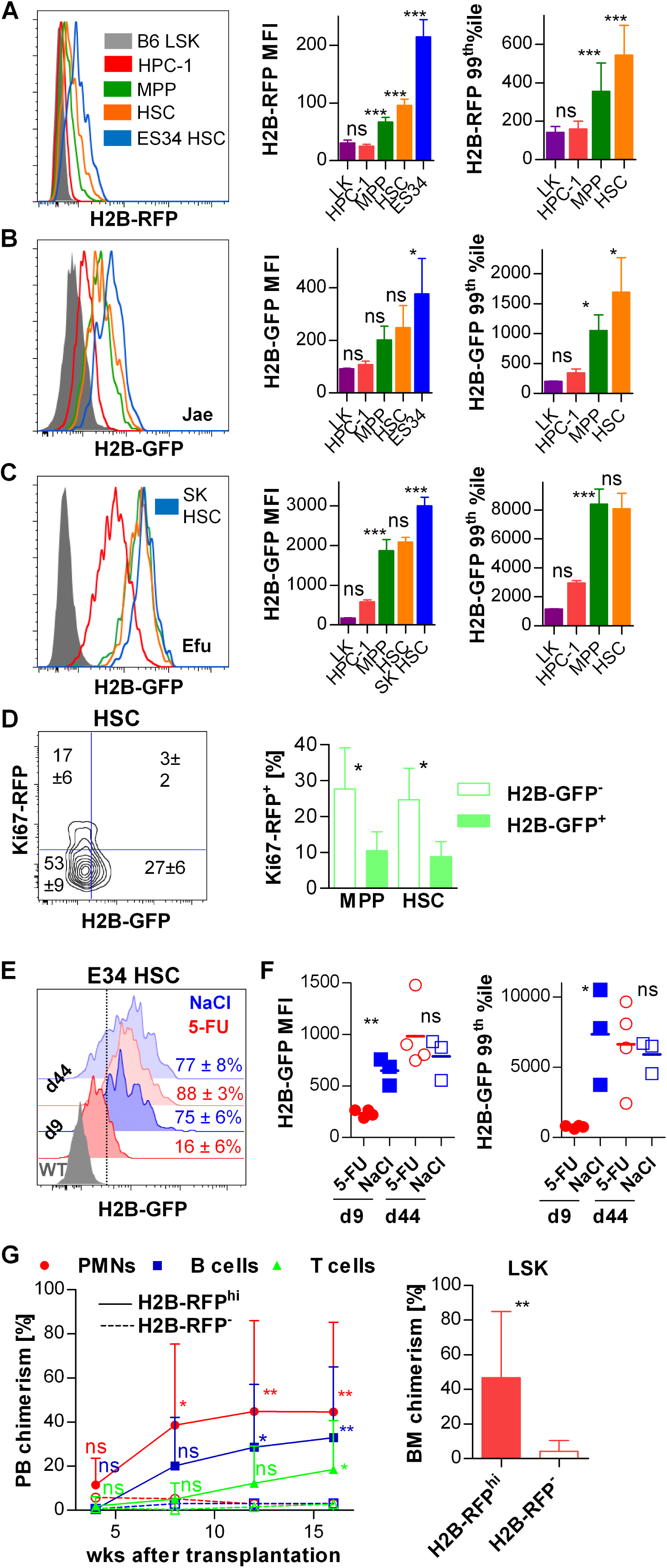
Primitive HSPCs exhibit higher levels of leaky H2B-FP background fluorescence. **A-C** BM HSPCs from repressed H2B-FP transgenic mouse models were analyzed for background fluorescence by flow cytometry. Histograms (left panels) show representative examples of H2B-FP background fluorescence of HSPC populations (LSK: lin^−/lo^Sca-1^+^Kit^+^; HPC-1: LSK CD48^hi^CD150^−^ (red); MPP: LSK CD48^−/lo^CD150^−^ (green); HSC: LSK CD48^−/lo^CD150^+^ (orange); ES34 HSC: lin^−/lo^Sca-1^hi^Kit^+^ CD48^−/lo^CD150^+^CD201^hi^CD34^−/lo^ (blue, Figure 1A-B); SK HSC: lin^−/lo^Sca-1^hi^Kit^lo^ CD48^−/lo^CD150^+^ (blue, Figure 1C); LK: lin^−/lo^Sca-1^−^Kit^+^ (purple); see Figure S1A for detailed gating strategy). LSK cells from WT B6 mice served as auto-fluorescence controls (grey tinted histogram). Median (MFI, middle data plot) and maximum (99^th^ percentile, right data plot) fluorescence intensity of BM HSPCs were determined. Significance was calculated by one-way repeated measures ANOVA with Bonferroni correction for multiple testing. **A** H2B-RFP background fluorescence of HSPCs isolated from un-induced *R26*^rtTA^/*Col1A1*^H2B-RFP^ mice (n=12, *Col1A1*^H2B-RFP/H2B-RFP^ (n=3) and *Col1A1*^H2B-RFP/wt^ (n=9, zygosity was corrected by doubling the fluorescence of heterozygous mice, see Figure S1B), 10-17 wks of age). **B** H2B-GFP background fluorescence of HSPCs isolated from un-induced *R26*^rtTA/rtTA^/*Col1A1*^H2B-GFP/H2B-GFP^ (Jae) mice (n=4, 22 wks of age). **C** H2B-GFP background fluorescence of HSPCs isolated from single transgenic tetO-H2B-GFP47Efu/J animals (n=3, homozygous, 10 wks of age). **D** HSPCs from un-induced *Ki67*^RFP/wt^*/R26*^rtTA/wt^*/Col1A1*^H2B-GFP/wt^ mice (n=6, age 12 - 15 wks) were analyzed for Ki67-RFP and H2B-GFP expression (left, representative contour plot of the HSC population, mean frequency ± SD of each quadrant is shown). Thresholds for Ki67-RFP and H2B-GFP gating were set according to either *Ki67*^RFP/wt^ or *R26*^rtTA/wt^*/Col1A1*^H2B-GFP/wt^ single positive animals (not shown). The frequency of proliferative Ki67-RFP^+^ cells within H2B-GFP^−^ or H2B-GFP^+^ MPPs and HSCs was calculated (right data plot, significance was analyzed by Wilcoxon’s signed-rank test). **E, F** Repressed *R26*^rtTA/rtTA^/*Col1A1*^H2B-GFP/H2B-GFP^ mice (11 - 16 wks of age) were injected with 5-fluorouracile (5-FU, shown in red, n=8) or 0.9% saline (blue, n=6) and BM was analyzed 9 or 44 d later by flow cytometry. (E) Representative histograms of H2B-GFP fluorescence in CD201^hi^CD34^−/lo^ LSK CD48^−/lo^CD150^+^ cells (E34 HSCs) are shown, H2B-GFP^+^ cells (mean frequency ± SD shown) were gated according to a 5-FU treated WT mouse (auto-fluorescence control, grey histogram). (F) H2B-GFP median (left) and maximum (99^th^ percentile, right) fluorescence intensity of E34 HSCs from 5-FU and saline-treated *R26*^rtTA/rtTA^/*Col1A1*^H2B-GFP/H2B-GFP^ individual mice are shown (significance was calculated by un-paired Student’s t test). **G** 50 H2B-RFP^−^ or H2B-RFP^hi^ HSCs were purified from un-induced *R26*^rtTA/wt^*/Col1A1*^H2B-RFP/wt^ animals (n=4, 9-12 wks, see Figure S1L for sorting strategy) and transplanted together with 3 x 10^5^ B6.CD45.1 WBMCs into irradiated B6.CD45.1/.2 recipient mice (n=8-9 / per group). PB (left plot) neutrophils (PMN, CD11b^+^Gr-1^hi^), B- (CD19^+^) and T-lymphocytes (CD3^+^) of chimeric recipients were analyzed for donor origin at indicated time points (mean and SD are shown, significance was calculated by repeated measures two-way ANOVA with Bonferroni post test). BM LSK chimerism was determined 17 weeks after transplantation (right plot, un-paired Student’s t test). An independent replicate of this experiment including secondary transplantation is shown in Figure S1M.

In order to prove that H2B-FP background fluorescence depends on mitotic activity, we forced quiescent HSCs of un-induced *R26*^rtTA/rtTA^/*Col1A1*^H2B-GFP/H2B-GFP^ mice into cell cycle by a single injection of 5-fluorouracil (5-FU). PB analysis 6 d later revealed uniform myeloablation in 5-FU-treated animals (Figure S1K). The H2B-GFP background fluorescence of BM CD201^hi^CD34^−/lo^ (E34) HSCs isolated from 5-FU injected mice 9 d after treatment was strongly reduced compared to saline-injected control mice (Figure 1E and 1F). A sub-cohort of animals was analyzed 44 d after 5-FU treatment and revealed that E34 HSCs had re-established high background fluorescence, similar to saline-treated control mice. This clearly demonstrated that enforced mitotic activity of HSCs led to temporary, cell cycle-dependent reduction of H2B background fluorescence.

As predominantly quiescent HSCs with a primitive immuno-phenotype exhibited bright H2B-FP background fluorescence, we reasoned that high background might identify HSCs with superior repopulating activity upon transplantation. Therefore, HSCs from un-induced *R26*^rtTA/wt^*/Col1A1*^H2B-RFP/wt^ donor mice were separated into H2B-RFP^−^ and H2B-RFP^hi^ populations (Figure S1L) and transplanted along with WT whole bone marrow competitor cells into lethally irradiated congenic recipient mice. PB and BM analysis of primary and secondary recipient mice (Figure 1G and S1M) revealed that H2B-RFP^−^ donor HSCs contained little long-term repopulation activity while transplantation of H2B-RFP^hi^ donor HSCs resulted in robust serial multi-lineage repopulation. Taken together, our experiments demonstrated that primitive HSCs accumulated high H2B-FP background fluorescence, while HSCs with dull background divided more frequently, up-regulated transcription- and translation-related genes and had significantly lower repopulation potential. Since all HSPC populations exhibited considerable background fluorescence and re-acquired background after temporary reduction in response to 5-FU, we conclude that there is continuous leaky background expression of H2B-FP even in the absence of label induction.

### Major contribution of H2B-FP degradation to loss of label

We observed much faster dilution of H2B-RFP (Morcos et al., 2017; Säwén et al., 2016) compared to H2B-GFP (Foudi et al., 2008; Oguro et al., 2013) in label retention experiments (Figure 2A) employing homologous Tet-controlled *R26*^rtTA^/*Col1A1*^H2B-FP^ transgenic systems (Beard et al., 2006) and suspected this to be caused by different stability of both H2B-FPs. In order to determine the rate of H2B-FP degradation, we assumed that, within short chase intervals, at least a fraction of HSCs displaying the highest H2B-FP retention will remain quiescent and will not or very rarely divide within this time period. Thus, over short chase intervals, the loss of fluorescence in those HSCs expressing the brightest H2B-FP fluorescence at each time point of analysis can be attributed to H2B-FP degradation. A similar approach in epithelial stem cells previously determined an H2B-GFP protein half-life of 24 d (Waghmare et al., 2008). Accordingly, we plotted a fluorescence time course of HSCs with maximum label retention isolated from pulsed and chased *R26*^rtTA^*/Col1A1*^H2B-RFP^ and *R26*^rtTA^*/Col1A1*^H2B-GFP^ mice (Figure 2B) and estimated protein degradation half-lives of around 2 and 6 weeks for H2B-RFP and H2B-GFP, respectively. To unambiguously prove the different stability of both H2B-FPs, we generated *R26*^rtTA^*/Col1A1*^H2B-GFP/H2B-RFP^ mice in which both labels are simultaneously expressed. A pulse-chase experiment employing these mice revealed much faster decay of H2B-RFP than H2B-GFP and verified previously estimated protein half-lives (Figure 2C-E). These rather short half-lives demonstrate that protein degradation is the major process responsible for loss of H2B-FP in quiescent HSCs. Therefore, neglecting this process must result in massive overestimation of HSC divisional activity.

**Figure 2.**
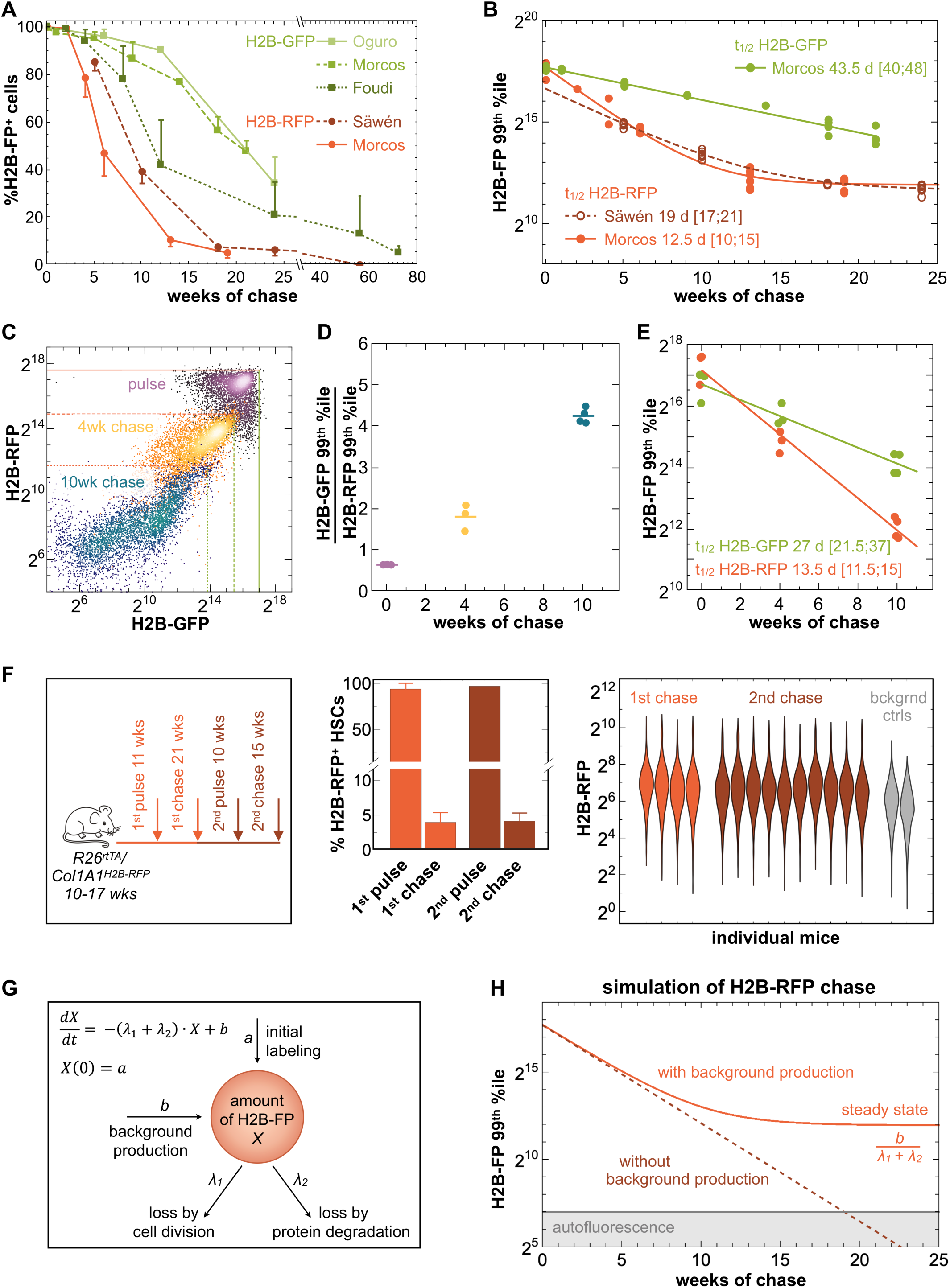
Major contribution of H2B-FP degradation to loss of label. **A** The frequencies of label-retaining (H2B-FP^+^) HSCs from pulsed and chased *R26*^rtTA^/*Col1A1*^H2B-FP^ mice are shown (H2B-RFP, red; H2B-GFP, green). Data was retrieved from Foudi et al., Nat Biotech 2008; Oguro et al., Cell Stem Cell 2013; Säwen et al., Cell Reports 2016 and Morcos et al., Stem Cell Reports 2017. H2B-GFP Morcos: *R26*^rtTA/rtTA^/*Col1A1*^H2B-GFP/H2B-GFP^ mice (n=1-4 / time point) were fed for 3 wks with DOX chow (2 g/kg) and chased for 0 - 21 wks. Mean frequencies and SD of LSK CD48^−/lo^CD150^+^ cells are shown, except for data from Oguro et al. in which a weighted mean of the HSC-1 and HSC-2 subpopulations was calculated ([1x HSC-1 + 3x HSC-2]/4). **B** Maximum H2B-FP fluorescence intensities (as judged by the 99^th^ percentile) of HSCs (LSK CD48^−/lo^CD150^+^) isolated from chased *R26*^rtTA^/*Col1A1*^H2B-RFP^ (shown in red; Säwen 2016 dashed line, open circles; Morcos 2017, continuous line, filled circles) or *R26*^rtTA/rtTA^/*Col1A1*^H2B-GFP/H2B-GFP^ (green; same animals as in Figure 2A ‘GFP Morcos’) individual mice are depicted. The data was fitted by our mathematical model (see Figure 2G) and loss of maximum fluorescence was attributed to H2B-FP degradation. H2B-FP half-lives and respective 95% confidence intervals (squared brackets) were calculated. **C-E** LSK CD48^−/lo^CD150^+^ HSCs of either pulsed (2g DOX /kg chow ad libitum for 7 wks), shown in purple), 4 (yellow), or 10 (blue) wk-chased *R26*^rtTA/wt^/*Col1A1*^H2B-GFP/H2B-RFP^ mice (n=3-4 / each time point) were analyzed for H2B-GFP and H2B-RFP fluorescence. **C** Representative examples of HSCs from either pulsed, 4 or 10 wk-chased *R26*^rtTA/wt^/*Col1A1*^H2B-GFP/H2B-RFP^ animals. Lines depict maximum (99^th^ %tile) H2B-GFP (shown in green) and H2B-RFP (red) fluorescence intensities of pulsed (continuous lines) and chased (dotted lines) mice. **D** The ratio of maximum H2B-GFP and H2B-RFP fluorescence intensity in HSCs was calculated for pulsed and chased animals (individuals and means are shown). **E** Time course of maximum H2B-GFP and H2B-RFP fluorescence of HSCs isolated from pulsed and chased *R26*^rtTA/wt^/*Col1A1*^H2B-GFP/H2B-RFP^ individuals. Calculation of H2B-FP half lives and respective 95% confidence intervals (squared brackets) was performed as in Figure 2B. **F** *R26*^rtTA/wt^*/Col1A1*^H2B-RFP/wt^ mice (n=21, 10-17 wks of age) were subjected to two consecutive cycles of H2B-RFP labeling by DOX induction (625 mg DOX / kg chow ad libitum for 10-11 wks, ‘pulse’) and dilution (‘chase’). Cohorts of animals were sacrificed at indicated time points for BM analysis (arrows) Frequencies of H2B-RFP^+^ HSCs (middle panel, LSK CD48^−/lo^CD150^+^) were analyzed after the 1^st^ (n=4 animals) and 2^nd^ (n=10) pulse-chase cycle. Violin plots (right panel) depict H2B-RFP fluorescence distributions of individual mice after the 1^st^ and 2^nd^ chase cycle. Un-induced *R26*^rtTA/wt^*/Col1A1*^H2B-RFP/wt^ mice (grey, 12 wks of age) served as background controls. **G** Overview of the ordinary differential equation model describing label dilution by accounting for 4 processes that affect average H2B-FP labeling intensity (*X*): (i) initial labeling *a* by H2B-FP induction, (ii) permanent leaky background H2B-FP production with constant rate *b*, (iii) H2B-FP decay due to division with rate λ*_1_*, and (iv) division-independent decay due to H2B-FP protein degradation with rate λ*_2_* (for details see methods). **H** Simulation of H2B-RFP label decay over time in pulsed and chased mice in the absence (dark red, dashed) or presence (red, continuous line) of continuous leaky background production. Grey area reflects auto-fluorescence.

As a gedankenexperiment, we neglect division-independent label degradation as implied by Bernitz et al. (2016) and further adhere to their conclusion that primitive HSCs undergo only 4 symmetric self-renewal divisions within the first 9 months of their life before they completely stop dividing. Under these assumptions, the H2B-FP dilution kinetics of young (<9 months) and aged (>9 months) mice should strongly differ from each other. In order to contest this prediction, we performed two consecutive cycles of doxycycline (DOX) induction and successive chase of *R26*^rtTA^/*Col1A1*^H2B-RFP^ mice (Figure 2F). We found similar labeling of HSCs after the first and second round of pulse. Cohorts of mice were analyzed after the first and second chase cycle and we observed highly similar label dilution in young and aged HSCs. If both label degradation had been absent and HSCs had entered permanent quiescence upon ageing, we would have observed a subpopulation of HSC retaining high levels of label after the second pulse. Competitive transplantation after the second pulse-chase cycle revealed an enrichment of repopulation activity among H2B-RFP^+^ cells, but also the H2B-RFP^−^ fraction contained HSCs with long term, multi-lineage potential (Figure S1N). Taken together, this experiment strongly argues against permanent quiescence in the absence of label degradation in ageing HSCs.

### Mathematical modeling to quantify label dilution kinetics

In order to quantify the impact of leaky background expression and division-independent degradation of H2B-FPs, we complemented the findings described above by a simple mathematical modeling approach accounting for key features of a pulse-chase experiment (Figure 2G). In particular, the model describes the kinetics of the average H2B-FP intensity *X* that is subject to four processes: (i) initial labeling *a* in case of H2B-FP induction, (ii) permanent leaky background H2B-FP production with constant rate *b*, (iii) H2B-FP decay due to division with rate λ*_1_*, and (iv) division-independent decay due to H2B-FP protein degradation with rate λ*_2_*.

In the absence of leaky background production, the model reduces to an exponential decay process, which does not reproduce the observed background expression above auto-fluorescence levels (Figure 2H). In contrast, assuming constant background label production, the model shows the pattern observed in the data: the decline of intensity is slowing down until reaching a constant background level. This final background level is characterized by a balance between the leaky background production rate *b* and the sum of the two decay processes λ*_1_* and λ*_2_*, but does not depend on the initial amount of label. This explains the observation of higher H2B-FP background fluorescence in populations with less proliferative activity: A lower division rate λ*_1_* results in a higher final H2B-FP accumulation.

In summary, the mathematical modeling strongly suggests that continuous leaky background expression is critical to explain the observed non-exponential dynamics of label dilution. We show that the final level of background fluorescence depends on both the degradation half-life of the FP and the mitotic activity of the cell population under investigation, but not on the initial amount of label in pulse-chase experiments.

### Age-dependent accumulation of H2B-FP background fluorescence

Based on our observation that quiescent HSPCs isolated from repressed H2B-FP mice exhibit high background fluorescence, we propose a scenario in which HSPC subpopulations start to accumulate H2B-FP background fluorescence upon entering extended phases of quiescence in early postnatal life (Bowie et al., 2006), while more proliferative cells will continue to constantly dilute background label via mitosis. In fact, mathematical modeling predicted that the accumulation of maximum H2B-FP background fluorescence would take weeks to several months after cessation of rapid cell divisions in early postnatal development (Figure 3A). The length of this time span depends on stability of H2B-FP as well as mitotic activity of the cell population under investigation. To rigorously test these predictions, we analyzed repressed mice of different ages from three H2B-FP transgenic mouse strains and found an age-dependent increase of median and maximum background fluorescence in various HSPC populations (Figure 3B-D). Our analysis revealed that (i) more quiescent HSPC populations took longer to build up final background fluorescence levels (e.g. HPC-1 < MPP < HSC) and (ii) the median and maximum background of HSPCs from all H2B-FP mouse models under investigation approached a plateau in older animals. These findings strongly confirmed the predictions derived from our mathematical model. A control analysis of differently aged WT B6 mice excluded any potential age-dependent increase of auto-fluorescence (Figure 3B-D).

**Figure 3.**
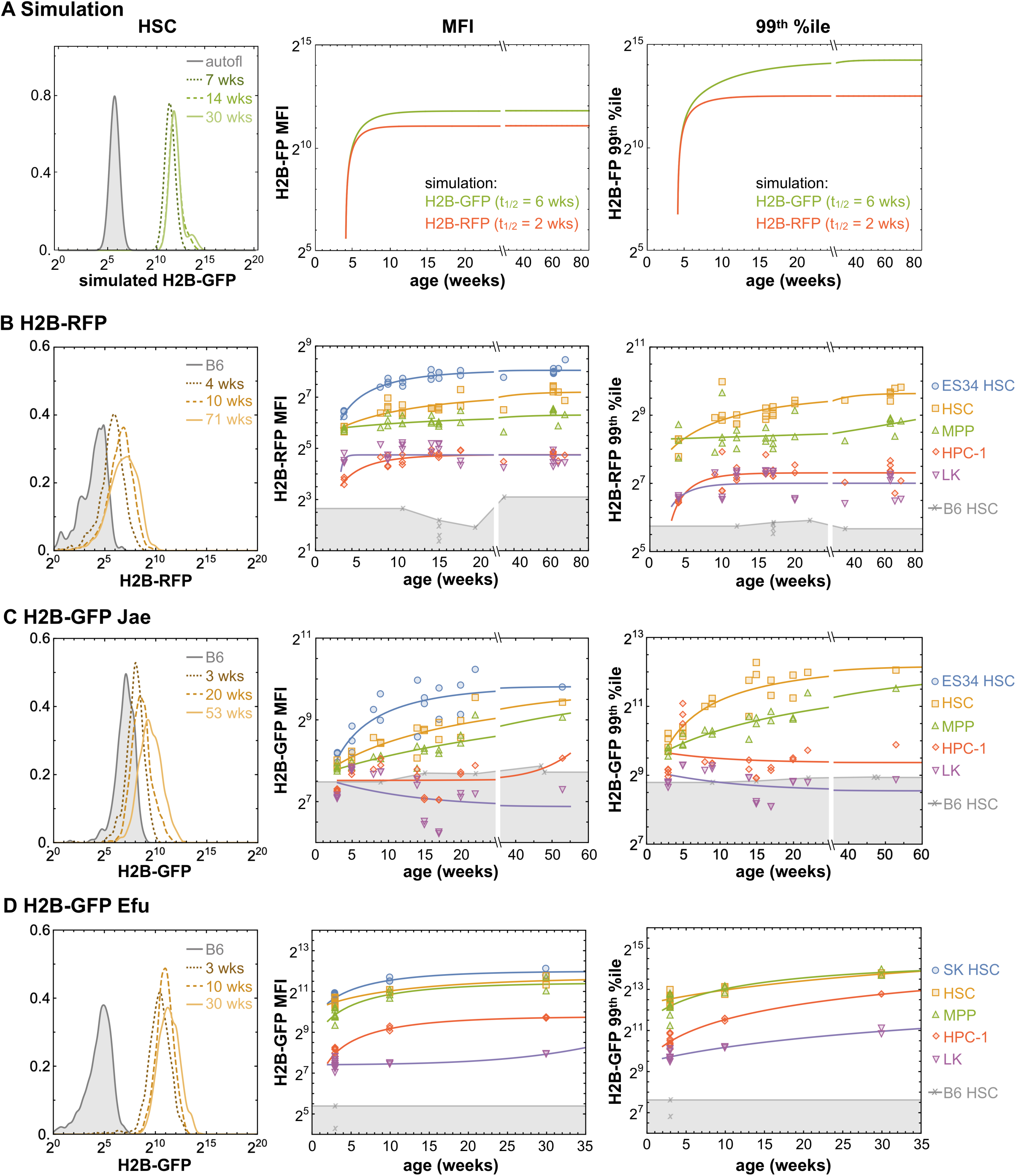
Age-dependent accumulation of H2B-FP background fluorescence. **A** Simulation of background labeling in repressed H2B-FP mouse models revealed accumulation of background fluorescence in predominantly quiescent HSCs upon ageing. Simulated H2B-GFP fluorescence distributions (left histograms) of HSCs from differently aged background controls are depicted. Median (MFI, middle panel) and maximum (99^th^ percentile, right panel) fluorescence intensities of HSCs expressing either H2B-GFP (shown in green, degradation t_1/2_= 6 wks) or H2B-RFP (red, degradation t_1/2_= 2 wks) background were simulated under the assumption that adult hematopoiesis is established after 4 weeks of age (see also methods). **B-D** BM HSPCs isolated from repressed H2B-FP mouse strains of various ages were analyzed for H2B-FP background fluorescence by flow cytometry. Representative examples of BM HSCs (LSK CD48^−/lo^CD150^+^) depict the age-dependent increase of H2B-FP background fluorescence (left histograms). Median (MFI, middle data plots) and maximum (99^th^ percentile, right data plots) H2B-FP fluorescence intensity of BM HSPCs were determined. Individual mice are shown; curves show fits of the mathematical model. HSCs from WT B6 mice (3-48 wks of age, grey crosses) served as auto-fluorescence controls. **B** H2B-RFP background fluorescence of HSPCs isolated from un-induced *R26*^rtTA^/*Col1A1*^H2B-RFP^ mice (n=25, *Col1A1*^H2B-RFP/H2B-RFP^ (n=11) and *Col1A1*^H2B-RFP/wt^ (n=14, zygosity was corrected by doubling the fluorescence of heterozygous mice). **C** H2B-GFP background fluorescence of HSPCs isolated from un-induced *R26*^rtTA/rtTA^/*Col1A1*^H2B-GFP/H2B-GFP^ mice (n=20). **D** H2B-GFP background fluorescence of HSPCs isolated from single transgenic tetO-H2B-GFP47Efu/J animals (n=13, homozygous: n=5, hemizygous: n=8, zygosity was corrected by doubling the fluorescence of hemizygous mice).

In summary, our data clearly demonstrates that H2B-FP background fluorescence of primitive HSCs progressively accumulated with age. Therefore, age-matched littermate background controls are crucial for accurate identification of H2B-FP-retaining HSCs in pulse-chase experiments.

### Mitotic activity of HSC populations inferred from H2B-FP label dilution data

We evaluated HSC label retention data by our mathematical model in order to disentangle the different sub-processes which account for loss of label (i.e. protein degradation, leaky background production and cell division). To resolve and quantify mitotic heterogeneity, we fitted different Gaussian distributions to the H2B-FP histograms of chased HSCs (Figure 4A and Figure S2A-B). For H2B-RFP-labeled HSCs, we identified two peaks (RFP^lo^ and RFP^hi^) throughout the entire chase (2 - 19 wks), while for chased HSCs expressing H2B-GFP, we initially found two peaks (GFP^lo^ and GFP^hi^), but starting from 14 wks of chase, a third, intermediate population (GFP^mid^) appeared in all replicates (Figure 4A-B).

**Figure 4.**
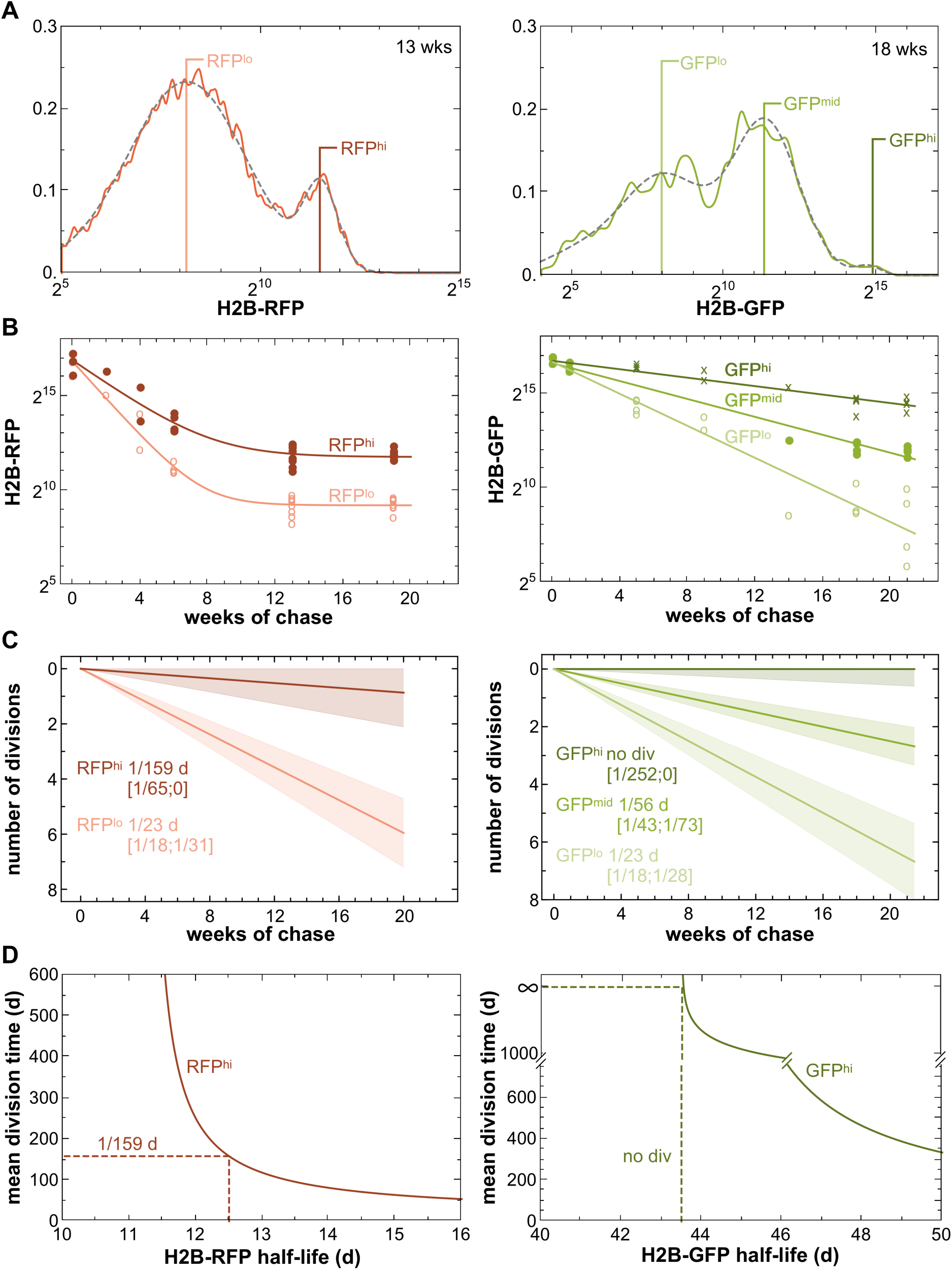
Mitotic activity of HSC populations inferred from H2B-FP label dilution data. **A** Representative examples for mixtures of Gaussian distributions fitted to H2B-RFP (left panel, data from Morcos 2017) or H2B-GFP (right panel, data from Figure 2A-B) label dilution data of HSCs (continuous curves: recorded data; dotted grey curves: fitted mixture distributions). Two Gaussians (RFP^lo^ and RFP^hi^, light and dark red, respectively) were fitted to H2B-RFP retention data (0-19 wks of chase), while for H2B-GFP retention data (0-21 wks of chase) two peaks (GFP^lo^ and GFP^hi^) were initially observed, but starting from 14 wks a third, intermediate peak uniformly appeared (GFP^mid^). The mean (horizontal lines) of each fitted Gaussian was estimated (see also Figure S2A-B). **B** H2B-FP dilution time courses of fitted Gaussians distributions for H2B-RFP (left; RFP^lo^: light red, open circles; RFP^hi^: dark red, filled circles) and H2B-GFP (right panel; GFP^lo^: light green, open circles; GFP^mid^: intermediate green, filled circles; GFP^hi^: dark green, crosses) HSC (LSK CD48^−/lo^CD150^+^) label retention data were plotted (curves show fits of the mathematical model, symbols depict means of fitted Gaussians from individual mice). **C** Division rates and their respective 95% confidence intervals (tinted areas and squared brackets) of RFP^lo^ and RFP^hi^ (left panel, red) or GFP^lo^, GFP^mid^ and GFP^hi^ (right panel, green) HSCs were calculated from data shown in B (for analysis of the CD34^−/lo^ HSC population see Figure S2C). **D** Model plots illustrating the dependence of the inferred mean division time within the most quiescent RFP^hi^ (left) or GFP^hi^ (right) HSC population on the accuracy of estimated H2B-FP degradation half-lives. Dotted lines depict the derived division rates for RFP^hi^ and GFP^hi^ HSCs based on the estimated H2B-RFP and H2B-GFP half-lives of 12.5 and 43.5 d, respectively.

For the most proliferative HSC subset, i.e. the cells which displayed the fastest loss of fluorescence (RFP^lo^ and GFP^lo^), we estimated an identical mean division rate of once per ~23 d from both H2B-RFP and H2B-GFP label dilution data sets (Figure 4C). This consistent division rate proved that the 3-fold difference in protein stability of H2B-GFP compared to H2B-RFP (t_1/2_ ~6 and 2 wks, respectively) accounted for their fundamentally different label retention characteristics (Figure 2A), while the genuine cell cycle activity of this HSC subset was not altered by the H2B-FP itself. Next, we analyzed H2B-FP dilution of predominantly quiescent HSC subpopulations, which retained higher levels of label. For the RFP^hi^ HSC population, we determined an average division rate of ~1/159 d. Employing our H2B-GFP data set, we estimated no or very rare divisions for the few GFP^hi^ HSCs, while the GFP^mid^ population, which appeared after 4 months of chase, displayed an average proliferation rate of 1/56 d. However, since protein degradation was the main reason for loss of label in quiescent HSCs, even in case of the more stable H2B-GFP, we found that our estimated division rates of predominantly quiescent RFP^hi^ and GFP^hi^ HSCs were highly imprecise as evidenced by their broad confidence intervals and a pronounced impact of the estimated H2B-FP protein degradation rate on the predicted division rate (Figure 4D). We conclude that the most quiescent HSCs of adult mice cycle infrequently (<1/100 d) and some may possibly never divide during the animal’s adulthood.

## Discussion

Our results demonstrate that loss of label in HSCs using different H2B-FPs is not only determined by label dilution via mitosis, but is additionally governed by division-independent degradation as well as leaky background production of label. We show that repressed H2B-FP mouse models progressively accumulate background fluorescence upon ageing to finally reach an equilibrium state in which loss of label and leaky background production are balanced. Consequently, high background fluorescence *per se* enriches for the most quiescent HSCs with a CD201^hi^, Sca-1^hi^ and CD34^−/lo^ immuno-phenotype and enhanced repopulation potential. We identified division-independent H2B-FP protein degradation as the major reason for loss of pulsed label in HSCs, precluding extended chase experiments and exact determination of mitotic history of quiescent HSCs. Nonetheless, our analysis of H2B-FP dilution data revealed infrequent, but steady mitosis of primitive adult HSCs.

### No evidence for mitotic memory and permanent quiescence of HSCs

Bernitz et al. (2016) proposed that a population of murine HSCs enters permanent quiescence after four symmetric self-renewal divisions within the first year of life. The authors hypothesized that these cells must count and remember their mitotic history by a yet unknown mechanism. This hypothesis is entirely based on the observation of a small HSC population with high repopulation potential seemingly retaining pulsed H2B-GFP fluorescence above background for up to 22 months of chase. While we disagree with their interpretation, the experimental results by Bernitz et al. (2016) are in good agreement with our own findings. Most prominently, Bernitz and colleagues consistently report a superior repopulation activity and increased quiescence of the HSC subpopulation with the highest levels of H2B-GFP fluorescence after long-term chase. However, their analysis does not account for continuous *de novo* H2B-GFP background production and its age-dependent and inevitable accumulation in quiescent HSC with high repopulation potential. Our data strongly suggests that quiescent HSCs with high background accumulation were mistaken for HSCs permanently retaining pulsed label, presumably by the use of inappropriate background controls (Figure S3A). The unambiguous identification of cells with minute levels of H2B-GFP retention, as reported by Bernitz et al. (2016), is further complicated by the considerable inter-individual variation of H2B-FP maximum background fluorescence within control mice of the same age (Figure 3 B-D).

Another serious concern regarding the validity of prolonged H2B-FP pulse-chase experiments is division-independent label degradation. For H2B-GFP, we determined a protein half-life of ~4-6 weeks in HSCs, which is similar to its stability in epithelial stem cells (Waghmare et al., 2008). Consequently, protein degradation on its own will reduce H2B-GFP fluorescence at least 500-fold, i.e. to undetectable levels, already after one year of chase (or >10^5^-fold after 22 months of chase) (Figure S3B). Our finding of substantial division-independent H2B-FP degradation is in line with the fact that various DNA transactions, like e.g. transcription, require histone displacement from DNA (Venkatesh and Workman, 2015). Even in non-replicating cells, high protein turnover was shown for endogenous H2B as well as for H2B-GFP (Jamai et al., 2007; Kimura and Cook, 2001; Toyama et al., 2013). Our pulse-chase experiment using mice with simultaneous expression of both H2B-FPs confirmed that the two fusion proteins differ in their stability. Moreover, two successive rounds of H2B-FP pulse-chase revealed unaltered label dilution in young and old mice (Figure 2F). This finding is not compatible with entry of a subpopulation of HSCs into permanent quiescence together with absence of H2B-FP degradation upon ageing.

Finally, we rebut counting of discrete HSC divisions using H2B-FP mouse models due to their inhomogeneous initial labeling. Bernitz et al. report an initial labeling distribution of ~2 decades of H2B-GFP fluorescence as well as considerable amounts of un-labeled HSCs (i.e. below background fluorescence) (Bernitz et al., 2016; Qiu et al., 2014). This might be explained by variable, HSC subset-specific activity of the hCD34 (Radomska et al., 2002) promoter driving H2B-GFP expression in this model. Use of the ubiquitous *R26*^rtTA^ Tet-driver with either *Col1A1*^H2B-FP^ allele resulted in a more homogenous initial H2B-FP distribution of HSCs with virtually 100% labeling above background and a peak width of ~ 1 decade (Foudi et al., 2008; Li et al., 2013; Nakada et al., 2014; Säwén et al., 2016) (Figure S3C). However, the broad initial HSC labeling range of ~1 - 2 decades of fluorescence intensity translates into 3 - 6 division equivalents already at start of chase. This broad heterogeneity renders counting of discrete HSC divisions impossible, which is in contrast to successful tracing of discrete divisions in homogeneously pulsed epithelial stem cells (Waghmare et al., 2008). A more homogenous labeling of a primitive HSC subpopulation that might be hidden within immuno-phenotypic HSCs was not detected (Figure S3D and S3E). Our modeling of H2B-FP retention revealed that loss of label slows down when cells approach the range of background fluorescence due to continuous *de novo* background production (Figure 2B, 2H and S3A-B). Therefore, mitosis of cells retaining pulsed label in the range of background fluorescence does not lead to halving of H2B-FP fluorescence and further obscures resolution of discrete division events within these cells.

### HSC mitotic activity in unperturbed adult bone marrow inferred from H2B-FP dilution

The rather short protein half-lives of H2B-FPs complicate the accurate estimation of mitotic activity in pulse-chase experiments as the major loss of label in primitive HSCs, which are predominantly quiescent, results from degradation rather than proliferation. Therefore, complete disentanglement of the respective contribution from either cell division or protein decay to loss of H2B-FP fluorescence in the most quiescent HSC subsets (i.e. RFP^hi^ or GFP^hi^) remained challenging (Figure 4D). However, our finding of infrequent mitotic activity among primitive HSCs is well compatible with the experimental notion that HSCs are not required for steady-state hematopoiesis in adult mice (Schoedel et al., 2016; Sheikh et al., 2016). This is further supported by fate mapping experiments (Busch et al., 2015; Sun et al., 2014), which reported rare contribution of HSCs to adult steady state hematopoiesis. Moreover, proliferation rates of primitive HSCs inferred from either BrdU uptake (1/55 d for HSC-1 (LSK CD48^−/lo^CD150^+^CD229^−/lo^CD244^−^) (Höfer et al., 2016; Oguro et al., 2013) or BrdU retention (1/145-193d) (van der Wath et al., 2009; Wilson et al., 2008) are in accordance with our conclusion that quiescent HSCs rarely but steadily divide as well as H2B-FP protein degradation being the major reason for loss of label over time in these cells. We consistently estimated a proliferation rate of 1/23 d in actively cycling (RFP^lo^ or GFP^lo^) LSK CD48^−/lo^CD150^+^ HSCs from both H2B-FP systems after correction for H2B-FP degradation, which is faster than the rate of 1/28-1/36 for ‘active’ HSCs inferred from BrdU retention (van der Wath et al., 2009; Wilson et al., 2008). However, the latter studies identified immuno-phenotypic HSCs by exclusion of CD34^+^ cells, a marker known to be upregulated in proliferating HSCs (Sato et al., 1999). Our analysis of H2B-RFP retention in LSK CD48^−/lo^CD150^+^CD34^−/lo^ cells revealed a division rate of 1/40 d for the proliferative RFP^lo^ subpopulation (Figure S2C), which is highly similar to the division rate of ‘active’ HSCs (van der Wath et al., 2009; Wilson et al., 2008) and substantiates our modeling approach. The division rates of proliferative HSCs (~1/8-1/18 d) and slowly cycling HSCs (~1/55–1/120 d) previously estimated from H2B-GFP retention data (Foudi et al., 2008) likely represent systematic overestimation due to neglect of label degradation, illustrating that consideration of label degradation is of particular importance in infrequently dividing cells.

In summary, H2B-FP transgenic models are important tools ideally suited to analyze and isolate viable cells based on differential mitotic activity. However, long-term tracing of discrete divisions of slowly cycling HSCs with label retention in the range of background fluorescence as performed by Bernitz et al. (2016) is not possible. The loss of label at early chase time points in these label retention experiments was mistakenly attributed to four discrete cell divisions, while our data strongly suggest that this loss of label largely represented protein degradation. At longer chase intervals, accumulation of high background fluorescence was most likely misinterpreted as stable retention of previously pulsed label by Bernitz et al (2016). Accounting for division-independent protein decay as well as for age-dependent background label accumulation is essential for estimating the divisional activity of rarely dividing cell populations in H2B-FP pulse-chase experiments. We found no evidence for abrupt entry into permanent quiescence upon ageing and neither for mitotic memory of HSCs. We rather conclude from our data that primitive HSCs continue to cycle rarely in aged mice.

## Author Contributions

M.N.F.M., C.M.M. and A.G. designed, performed, and analyzed experiments. M.N.F.M., A.G., T.Z. and I.G. conceptualized data analysis. T.Z and I.G. conceived mathematical modeling. T.Z. performed quantitative data analysis and implemented the mathematical modeling. Y.G., A.P., S.R. and A.D. performed transcriptome analysis. P.S., H.W., D.B., N.A., R.B. and M.M. provided H2B-FP data. A.G. conceived and supervised the study and wrote the manuscript. A.R. advised the study and edited the manuscript. All authors read, commented and agreed on the final version of the manuscript.

## Acknowledgements

We thank Livia Schulze, Tobias Häring, Christina Hiller and Christa Haase for technical assistance, Jeffrey Bernitz for scientific discussions, Thomas Höfer and Kristina Schödel for critical reading of the manuscript.

A.G. was supported by the German Research Council (DFG, GE3038/1-1). Y.G. and A.R. were funded by the German Research Council (RO2133/10-1) in the setting of FOR2577. M.N.F.M. was supported by the Dresden International Graduate School for Biomedicine and Bioengineering (DIGS-BB), granted by the German Research Council in the context of the Excellence Initiative. The work of T.Z. and I.G. was supported by the German Federal Ministry of Education and Research, grant 031A315 “MessAge” to I.G. P.S., H.W. and D.B. acknowledge funding from the Tobias Foundation and the Swedish Cancer Society.

The authors declare no competing financial interests.

## Materials and methods

### Experimental model and subject details

#### Mice

*R26*^rtTA^*/Col1A1*^H2B-RFP^ (Jax No: 014602) (Egli et al., 2007) and *R26*^rtTA^*/Col1A1*^H2B-GFP^ (Jax No: 016836) (Foudi et al., 2008), C57Bl/6JRj wt (Janvier), B6.CD45.1 (Jax No. 002014), B6CD45.1/.2, and Ki67-RFP (obtained from Clevers lab) (Basak et al., 2014) mice were housed at the Experimental Center, TU Dresden. Tg(tetO-HIST1H2BJ/GFP)47Efu/J (tetO-H2B-GFP47Efu/J) were imported from Jackson labs (Jax No. 005104) and housed at the animal facility of BMC Lund University. Tg(tetO-HIST1H2BJ/GFP)47Efu mice were backcrossed to C57Bl/6 genetic background for more than 10 generations to yield the B6.tetO-H2B-GFP47Efu strain, which was housed at DKFZ Heidelberg. Pulse-chase data of *R26*^rtTA^*/Col1A1*^H2B-RFP^ mice was previously published (Morcos et al., 2017; Säwén et al., 2016).

H2B-FP mice were induced with doxycyclin (DOX) either via chow (Ssniff Spezialdiäten, 625mg/kg for 10-11 wks or 2000 mg/kg for either 3-7 weeks (this study) or 1 week (Säwén et al., 2016)) or via drinking water (1 mg/ml DOX (Applichem), 1% sucrose for 8 weeks (Morcos et al., 2017)) ad libitum.

5-FU (150 µg/g body weight, Applichem) was administered via intravenous (i.v.) injection.

All animal experiments were in accordance with institutional guide lines and were approved by the relevant authorities (Landesdirektion Dresden, Regierungspräsidium Karlsruhe and local ethics committee University Lund).

### Methods Details

#### Cell Preparation

Whole bone marrow cells (WBMCs) were isolated by crushing long bones with mortar and pestle using PBS/2% FCS/2 mM EDTA and filtered through a 100 µm mesh. After erythrocyte lysis in hypotonic NH_4_Cl-buffer, cells were filtered through a 40 µm mesh. Hematopoietic lineage^+^ cells were removed with the lineage cell depletion kit (Miltenyi Biotec).

Peripheral blood (PB) was drawn into glass capillaries by retrobulbar puncture. For chimerism analysis, blood was flushed out of the capillary using PBS / Heparin (250u/ml, Biochrom). Erythrocyte lysis in NH_4_Cl-buffer was performed twice for 5 minutes. For hemograms, blood was drawn by retrobulbar puncture directly into EDTA-coated tubes (Sarstedt) and analyzed on a XT-2000i Vet analyzer (Sysmex).

#### BM transplantation

B6.CD45.1/.2 recipient mice received a single dose of 9 Gy total body irradiation (Yxlon Maxi Shot γ source). Donor cells (purified LSK CD48^−/lo^CD150^+^ test cells mixed with B6.CD45.1 competitor WBMCs) were administered via intravenous injection into the retro-orbital sinus. For secondary transplantation equal numbers of WBMCs from all primary transplanted individuals were pooled and 4 x 10^6^ cells were transplanted into each lethally irradiated secondary recipients. Recipient peripheral blood (PB) T-lymphocytes (CD3^+^), B-lymphocytes (B220^+^) and neutrophils (CD11b^+^, Gr-1^hi^) were analyzed for their donor origin using a MACSquant flow cytometer (Miltenyi).

#### Flow cytometry

Cells were incubated with antibodies (Table S1) in PBS / 2% FCS for 30 min, washed twice with PBS / 2% FCS and analyzed on a ARIA II SORP or ARIA III (BD Biosciences, Heidelberg, Germany). H2B-FP fluorescence intensities from independent experiments on the same flow cytometer were normalized using Sphero Rainbow 8 peak particles (BD Bioscience) as reference. Data were analyzed with FlowJo V9.9 software (Tree Star,) and gates were set according to Fluorescence-Minus-One (FMO) controls. For a detailed overview of gating strategies refer to Figure S1A.

#### Single cell RNAsequencing

Single BM LSK CD48^−/lo^CD150^+^ cells were index-sorted into 96 well plates containing 2 µl of nuclease free water with 0.2% Triton-X 100 and 4 U murine RNase Inhibitor (NEB), spun down and frozen at −80°C. After thawing, 2 µl of a primer mix were added (5 mM dNTP (Invitrogen), 0.5 µM dT-primer (C6-aminolinker-AAGCAGTGGTATCAACGCAGAGTCGACTTTTTTTTTTTTTTTTTTTTTTTTTTTTTTVN), 4 U RNase Inhibitor (NEB)). RNA was denatured for 3 minutes at 72°C and the reverse transcription (RT) was performed at 42°C for 90 min after filling up to 10 µl with RT buffer mix for a final concentration of 1x superscript II buffer (Invitrogen), 1 M betaine, 5 mM DTT, 6 mM MgCl_2_, 1 µM TSO-primer (AAGCAGTGGTATCAACGCAGAGTACATrGrGrG), 9 U RNase Inhibitor and 90 U Superscript II (Invitrogen). After synthesis, the reverse transcriptase was inactivated at 70°C for 15 min. The cDNA was amplified using Kapa HiFi HotStart Readymix (Peqlab) at a final 1x concentration and 0.1 µM UP-primer (AAGCAGTGGTATCAACGCAGAGT) under following cycling conditions: initial denaturation at 98°C for 3 min, 22 cycles [98°C 20 sec, 67°C 15 sec, 72°C 6 min] and final elongation at 72°C for 5 min. The amplified cDNA was purified using 1x volume of hydrophobic Sera-Mag SpeedBeads (GE Healthcare) and DNA was eluted in 12 µl nuclease free water. The concentration of the samples was measured with a Tecan plate reader Infinite 200 pro in 384 well black flat bottom low volume plates (Corning) using AccuBlue Broad range chemistry (Biotium).

For library preparation 700 pg cDNA in 2 µl was mixed with 0.5 µl Tagment DNA Enzyme and 2.5 µl Tagment DNA Buffer (Nextera, Illumina) and tagmented at 55°C for 5 min. Subsequently, Illumina indices were added during PCR (72°C 3 min, 98°C 30 sec, 12 cycles [98°C 10 sec, 63°C 20 sec, 72°C 1 min], 72°C 5 min) with 1x concentrated KAPA HiFi HotStart Ready Mix and 0.7 µM dual indexing primers. After PCR, libraries were quantified with AccuBlue Broad range chemistry, equimolarly pooled and purified twice with 1x volume Sera-Mag SpeedBeads. This was followed by Illumina single-end sequencing (76bp) on a Nextseq550 aiming at an average sequencing depth of 0.5 mio reads per cell.

#### Single cell transcriptome analysis

Raw reads were mapped to the mouse genome (mm10) and splice-site information from Ensembl release 87 (Zerbino et al., 2017) with gsnap (version 2018-07-04) (Nacu and Wu, 2010). Uniquely mapped reads and gene annotations from Ensembl were used as input for featureCounts (version1.6.2) (Smyth et al., 2014) to create counts per gene and cell.

Further analysis of the single cell counts data was done with the R package scater (version1.8.4) (McCarthy et al., 2017). Cells which met one of the following criteria were considered low quality: 1) total counts < 5,000, 2) total genes <2,000 or >10,000, 3) percentage of counts from mitochondrial genes > 6%, 4) percentage of counts from ERCC spike-in transcripts > 25% and 5) percentage of genes without counts >95%. Consequently, 82 low-quality cells (29%) were removed. Also, genes which were expressed in less than 1/3 of the remaining 200 cells were considered as low expressed and removed leaving 6,358 genes for further analysis.

For the filtered counts matrix (6,358 genes x 200 cells), counts per million (CPM) normalization was applied to alleviate the effect of different sequencing depth across the cells. To make expression value comparable, expression values of a gene were transformed to the scale [0,1] where 0 was the minimum and 1 the maximum CPM value of this gene across cells.

MoIO and NoMO genes (Wilson et al., 2015), which showed expression in our data, were considered (18 and 15 genes, respectively). The scores were then calculated by averaging the normalized scaled expression values of MoIO and NoMO gene sets, respectively. The relationship between fluorescence markers (H2B-GFP and Sca-1, in logarithmic scale), and MoIO/NoMO scores was calculated using linear regression.

To identify functional changes in gene expression correlated with H2B-GFP marker expression, for each gene its Spearman correlation coefficient to the fluorescence marker (H2B-GFP and Sca-1) was calculated. The top 100 positive and top 100 negative correlated genes were analyzed for associated terms from the Gene Ontology (GO) database, aspect: Biological Process (BP) using DAVID 6.8 (Huang et al., 2008). Very broad and unspecific GO BP terms were excluded by the FAT filter and similar/redundant terms were grouped into annotation clusters and enrichment scores were calculated. All clusters with an enrichment score higher than 1 were considered.

#### Mathematical modeling

The dilution kinetics of the fluorescent histone 2B fusion protein (H2B-FP) is formulated in terms of an ordinary differential equation describing the average kinetics of the amount of label, termed *X*, in a homogeneous population:

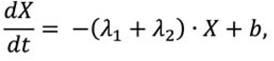

with (i) the rate of permanent leaky background FP production *b*, (ii) FP decay due to division with a rate λ*_1_*, and (iii) division-independent FP decay due to protein degradation with a rate λ*_2_*. The initial condition *X(0)=a* describes the induced amount of FP label *a* in case of an H2B-FP induction. This model assumes a homogeneous population constant in size. This means that cell differentiation balances the increase by cell division and differentiating cells are no longer considered as part of the modeled HSC population. Cell cycle times are assumed to follow an exponential distribution with mean 1/λ*_1_*. There is no memory or heritability in the process: The probability of a cell to divide does not depend on former divisions. Furthermore, the probability of differentiation (and subsequent extinction from the stem cell compartment) does not depend on the number of cell divisions that a cell has undergone.

The equation can be solved explicitly to

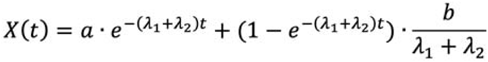

and was fitted to the logarithmic data using the nonlinear model fit routine in Mathematica (version 11.3, Wolfram Inc.).

In the long run the system reaches a steady state (i.e. a protein level that does not change over time) at b/(λ_1_+λ_2_), independent of initially provided amount of label *a*. At this equilibrium level, leaky background production (described by *b*) and decay due to cell division and protein degradation (described by λ*_1_*+λ*_2_*) are leveling out.

To determine the time until chased and background control populations merge (Figure S3B), we assume that the chase period starts at 10 weeks after birth, while the background accumulation is assumed to start at 4 weeks after birth when the adult hematopoiesis is established (Bowie et al., 2006), thus leading to a time shift between both populations of 6 weeks (=42 days). We define the time point of merging as the time when the chased population reaches at least 95% of the background control population. Assuming the limiting case of no cell division this leads to the following equation:

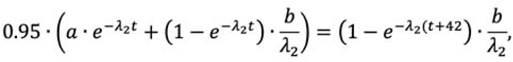

which resolves to:

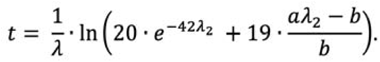

To study the process of establishing a stable level of H2B-FP background after birth (Figure 3A), a model comprised of two independent HSC populations was used based on the following assumptions: (i) The initial level of label at the time point of 4 weeks is assumed to be autofluorescence. This assumes that juvenile HSC cycle fast and thus do not or neglectably accumulate H2B-FP label until the hematopoietic system is established; (ii) we assume a mixture of two HSC populations: a proliferative population (90%) with an average cycling of 16 days and a quiescent population (10%) that does not cycle; (iii) for H2B-FP degradation, we assume a half-life of 2 weeks for H2B-RFP and 6 weeks for H2B-GFP as estimated.

To estimate subpopulations from H2B-FP distributions (Figure 4), a mixture distribution of up to four normal distributions was fitted to the logarithmic data. Estimated subpopulation means were then separately fitted by the mathematical model.

### Quantification and statistical analyses

Data are presented as mean and SD. Statistical analysis was done with Prism 5 (Graphpad), applied statistical tests are denominated in the respective figure legend. Significant results are indicated by: *p = 0.01–0.05, **p = 0.001–0.01, and ***p < 0.001; not significant results by: ns.

## Online supplemental material

Supplemental material includes three figures and one table. Figure S1 presents additional data on H2B-FP background accumulation in primitive HSPCs. Figure S2 provides representative examples of H2B-FP dilution after different chase intervals. Figure S3 shows simulations of H2B-FP pulse-chase experiments and data on labeling homogeneity. Table S1 lists antibody conjugates used for flow cytometry.

**Figure S1.**
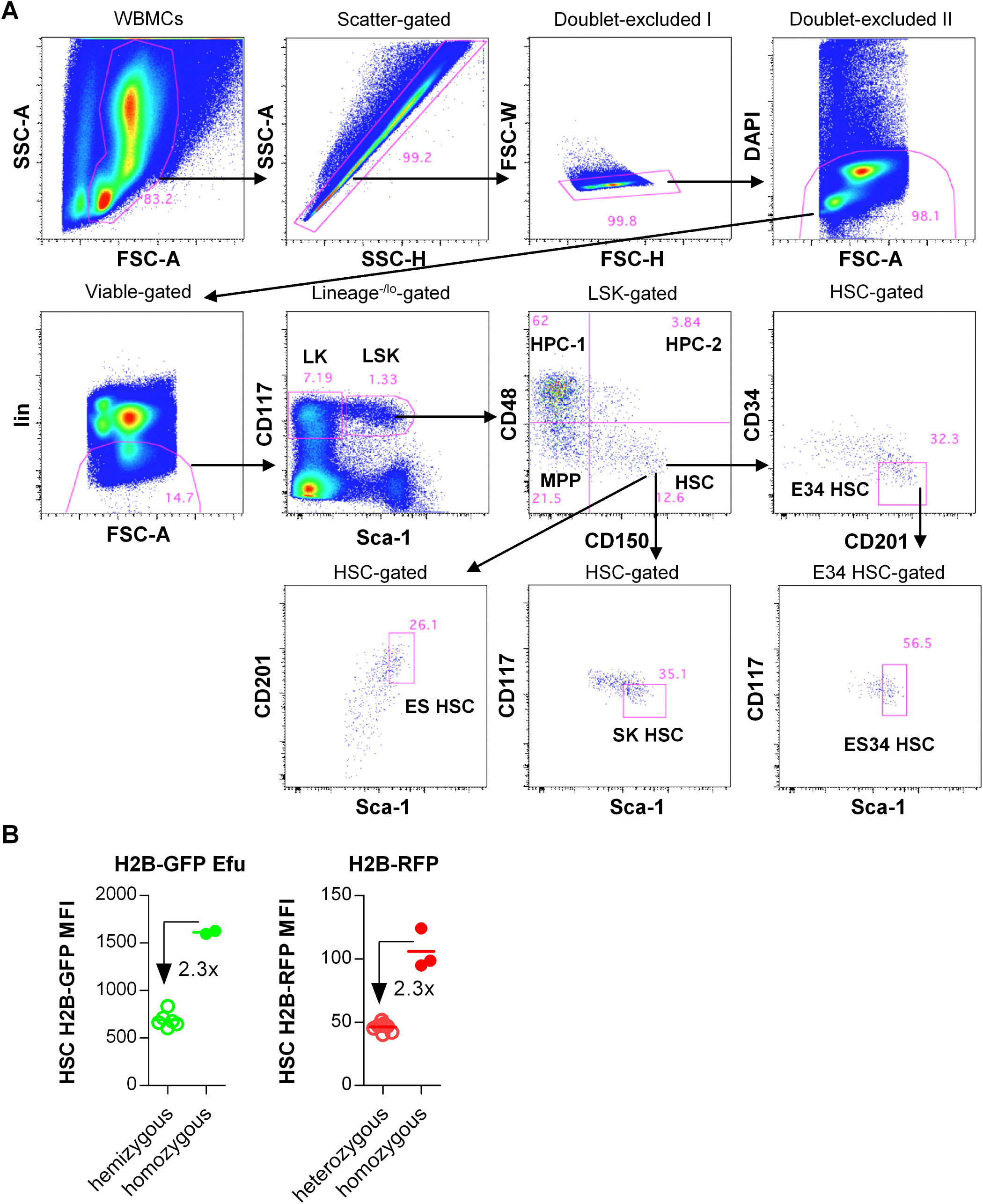
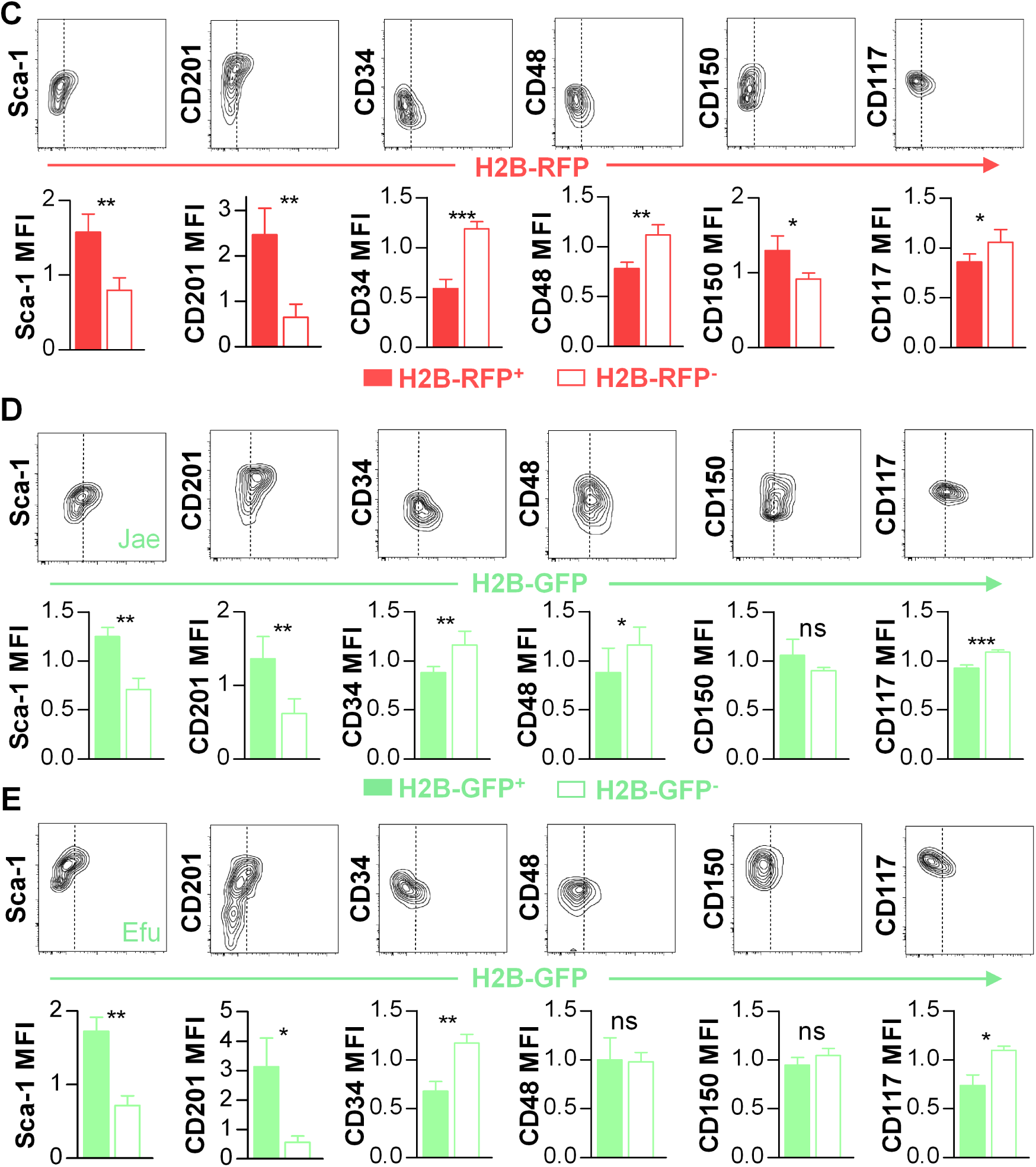
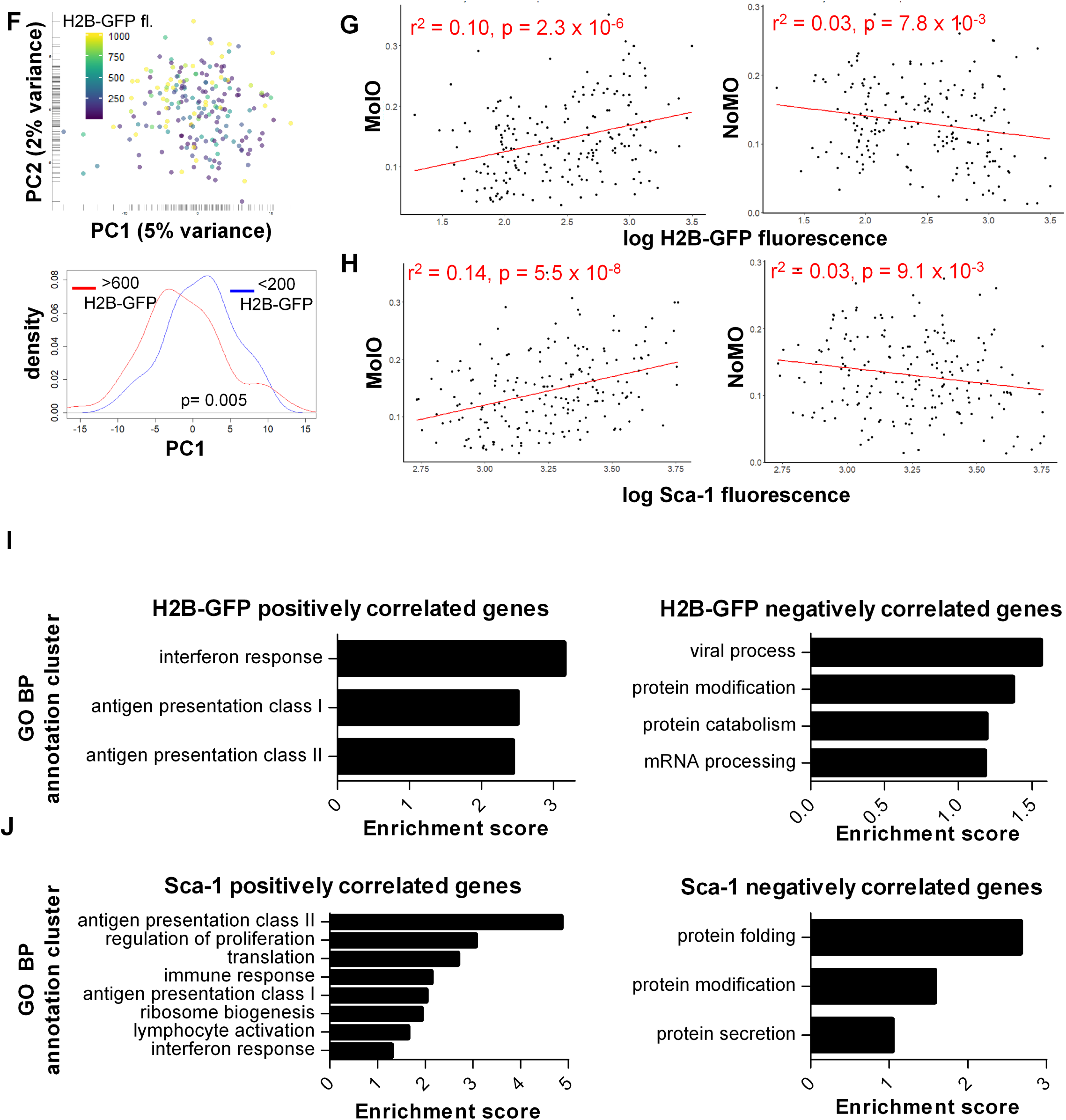
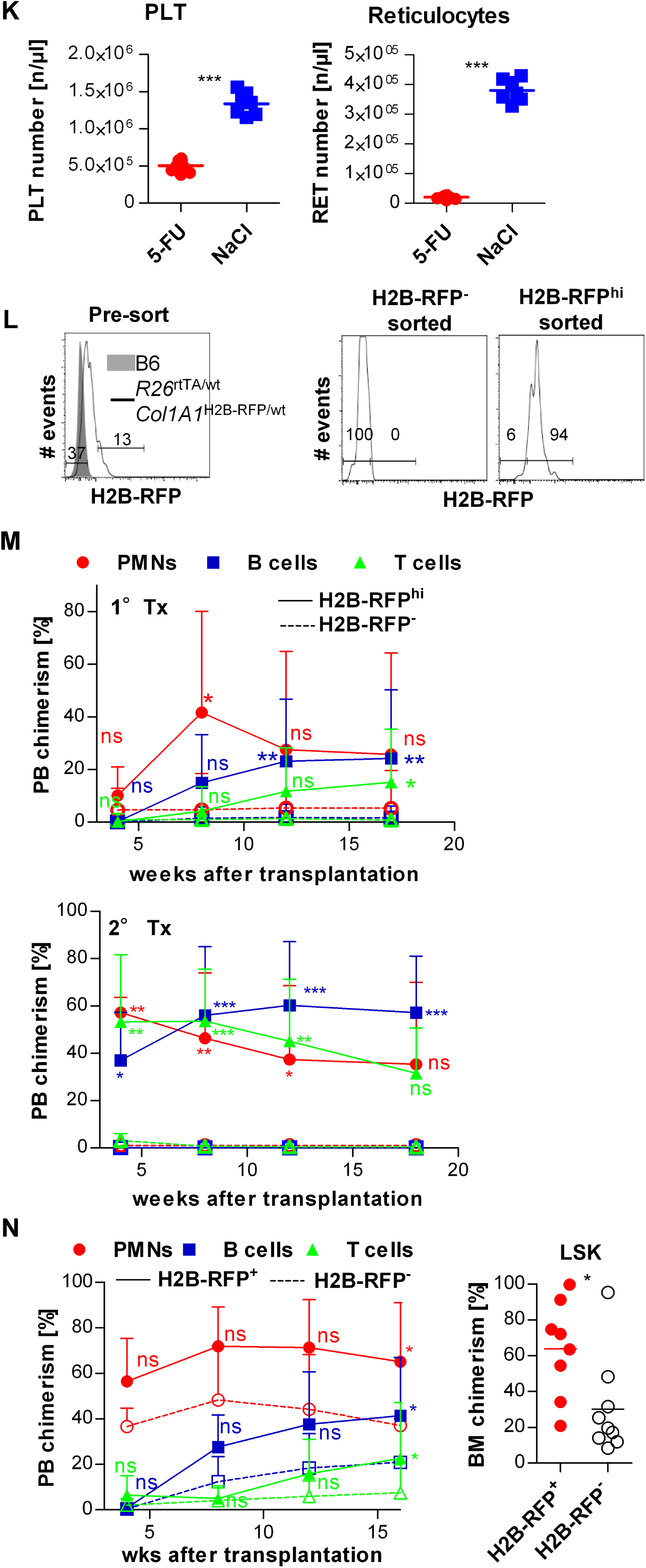
Primitive HSPCs exhibit higher levels of leaky H2B-FP background fluorescence. **A** Representative gating strategy of B6 WT HSPCs (LK: lin^−/lo^Sca-1^−^CD117^+^; LSK: lin^−/lo^Sca-1^+^CD117^+^; HPC-1: LSK CD48^hi^CD150^−^; HPC-2: LSK CD48^hi^CD150^+^; MPP: LSK CD48^−/lo^CD150^−^; HSC: LSK CD48^−/lo^CD150^+^; E34 HSC: LSK CD48^−/lo^CD150^+^CD34^−/lo^CD201^hi^; ES34 HSC: LS^hi^K CD48^−/lo^CD150^+^CD34^−/lo^CD201^hi^; SK HSC: LS^hi^K^lo^ CD48^−/lo^CD150^+^; ES HSC: LS^hi^K CD48^−/lo^CD150^+^CD201^hi^; FSC: forward scatter; SSC: side scatter; DAPI: 4′:6-diamidino-2-phenylindole; lin: hematopoietic lineage antigens). **B** Age-matched hemi- (n=6) or homozygous (n=2) single transgenic tetO-H2B-GFP47Efu/J as well as un-induced *R26*^rtTA^/*Col1A1*^H2B-RFP/wt^ (n=9, heterozygous) or *R26*^rtTA^/*Col1A1*^H2B-RFP/H2B-RFP^ (n=3, homozygous) mice were analyzed for H2B-FP MFI of HSCs (individuals and means (bars) are shown, fold of fluorescence reduction was calculated). **C-E** BM HSCs from repressed H2B-FPs transgenic mouse models were analyzed for background fluorescence and surface marker expression by flow cytometry. H2B-FP fluorescence of HSCs (LSK CD48^−/lo^CD150^+^) in relation to surface markers is depicted (representative contour plots upper rows, dotted lines show gating threshold for events which are within (H2B-FP^−^) or above (H2B-FP^+^) auto-fluorescence). Lower rows: Comparison of surface marker expression between H2B-FP^−^ and H2B-FP^+^ HSCs (means and SD are shown, each surface marker expression was normalized to the respective mean MFI of the total HSC population). Significance was calculated by paired Student’s t test. **C** BM HSCs of un-induced *R26*^rtTA/wt^/*Col1A1*^H2B-RFP/wt^ mice (n= 4, aged 9-12 wks) mice were analyzed for H2B-RFP background fluorescence and surface marker expression. **D** BM HSCs of un-induced R26^rtTA/rtTA^/*Col1A1*^H2B-GFP/H2B-GFP^ mice (n=4, age 22 wks) mice were analyzed for H2B-GFP background fluorescence and surface marker expression. **E** BM HSCs of hemizygous, single transgenic B6.tetO-H2B-GFP47Efu mice (n= 3, age 11-15 wks) were analyzed for H2B-GFP background fluorescence and surface marker expression. **F** Principal component (PC) analysis of 200 index-sorted single HSCs (LSK CD48^−/lo^CD150^+^) isolated from un-induced *R26*^rtTA/rtTA^/*Col1A1*^H2B-GFP/H2B-GFP^ mice (n= 6, aged 12 – 22 wks, data derived from two independent experiments) revealed separation of cells with high H2B-GFP background fluorescence (upper panel). Each cell was colored according to its H2B-GFP background fluorescence intensity (H2B-GFP fl.), which was recorded during index-sorting. PC1 distributions of cells (lower panel) expressing either > 600 (red histogram) or < 200 (blue histogram) units of H2B-GFP fluorescence reveal significant (Mann–Whitney U test) enrichment of cells with high H2B-GFP background fluorescence in the left area of the PCA plot. **G, H** The recorded H2B-GFP (G) or Sca-1 (H) fluorescence of each single cell was correlated to normalized gene signatures of quiescent (MolO, Wilson et al., Cell Stem Cell 2015) or proliferative (NoMO) HSCs and revealed a significant positive correlation of H2B-GFP background fluorescence as well as Sca-1 surface expression to the MolO score and concordant negative correlations to the NoMO score (r^2^ and significance was calculated by linear regression and F-test, respectively). The previously published (Wilson et al., 2015) correlations between the surface marker Sca-1 to the MolO and NoMO signatures had similar direction, strength and significance in our data set as the correlation of H2B-GFP background fluorescence to these gene signatures. **I, J** Plots show biological processes (BP) that are enriched in the first 100 genes positively (left panel) or negatively (right panel) correlated to either H2B-GFP background fluorescence (I) or Sca-1 surface expression (J). TOP100-negatively correlated genes were enriched for genes related to transcription and translation. Analysis was performed using the gene ontology (GO) BP database of DAVID 6.8. Similar terms were clustered and enrichment scores (>1, log transformation of the DAVID Expression Analysis Systematic Explorer [EASE] score) were calculated to determine overrepresentation of particular biological processes. **K** PB platelet (PLT) and reticulocyte (RET) counts of repressed *R26*^rtTA/rtTA^/*Col1A1*^H2B-GFP/H2B-GFP^ mice were determined 6 d after 5-FU or saline injection (NaCl). **L** Sorting strategy (left histogram) for isolation and re-analysis (right histograms) of H2B-RFP^−^ and H2B-RFP^hi^ HSCs (LSK CD48^−/lo^CD150^+^) from un-induced *R26*^rtTA/wt^*/Col1A1*^H2B-RFP/wt^ donor mice (n=4) is depicted. H2B-RFP^−^ gate was set according to the red auto-fluorescence of B6 WT HSCs. The event frequency in each gate is shown. **M** 30 H2B-RFP^−^ or H2-RFP^hi^ HSCs were purified from un-induced *R26*^rtTA/wt^*/Col1A1*^H2B-RFP/wt^ animals (n=6, 16-17 wks) and transplanted together with 2 x 10^5^ B6.CD45.1 WBMCs into irradiated B6.CD45.1/.2 recipient mice (n= 9-10 / per group). PB neutrophils (PMN, CD11b^+^Gr-1^hi^), B-(CD19^+^) and T-lymphocytes (CD3^+^) of primary recipients were analyzed for donor origin at indicated time points (upper panel). 4 x 10^6^ WBMCs were secondary transplanted 17 weeks after primary transplantation. PB neutrophils (PMN, CD11b^+^Gr-1^hi^), B-(CD19^+^) and T-lymphocytes (CD3^+^) of secondary recipients (n=4 / group) were analyzed for donor origin at indicated time points (lower panel, mean and SD are shown, significance was calculated by repeated measures two-way ANOVA with Bonferroni post test). **N** Competitive transplantation of *R26*^rtTA/wt^*/Col1A1*^H2B-RFP/wt^ animals after the 2^nd^ chase (Figure 2F). 200 H2B-RFP^+^ (continuous lines) or H2B-RFP^−^ (dotted lines) HSCs were purified, mixed with 3×10^5^ B6.CD45.1 WBMCs, and each transplanted into lethally irradiated wt recipients (n= 8-9 / donor cell type). PB neutrophil, B and T cell chimerism was analyzed for 16 weeks (left, significance was calculated by repeated measures two-way ANOVA with Bonferroni post test). BM LSK chimerism was analyzed after 17 wks (right panel, individual recipient mice and means are shown, significance was calculated by an un-paired Student’s t test).

**Figure S2.**
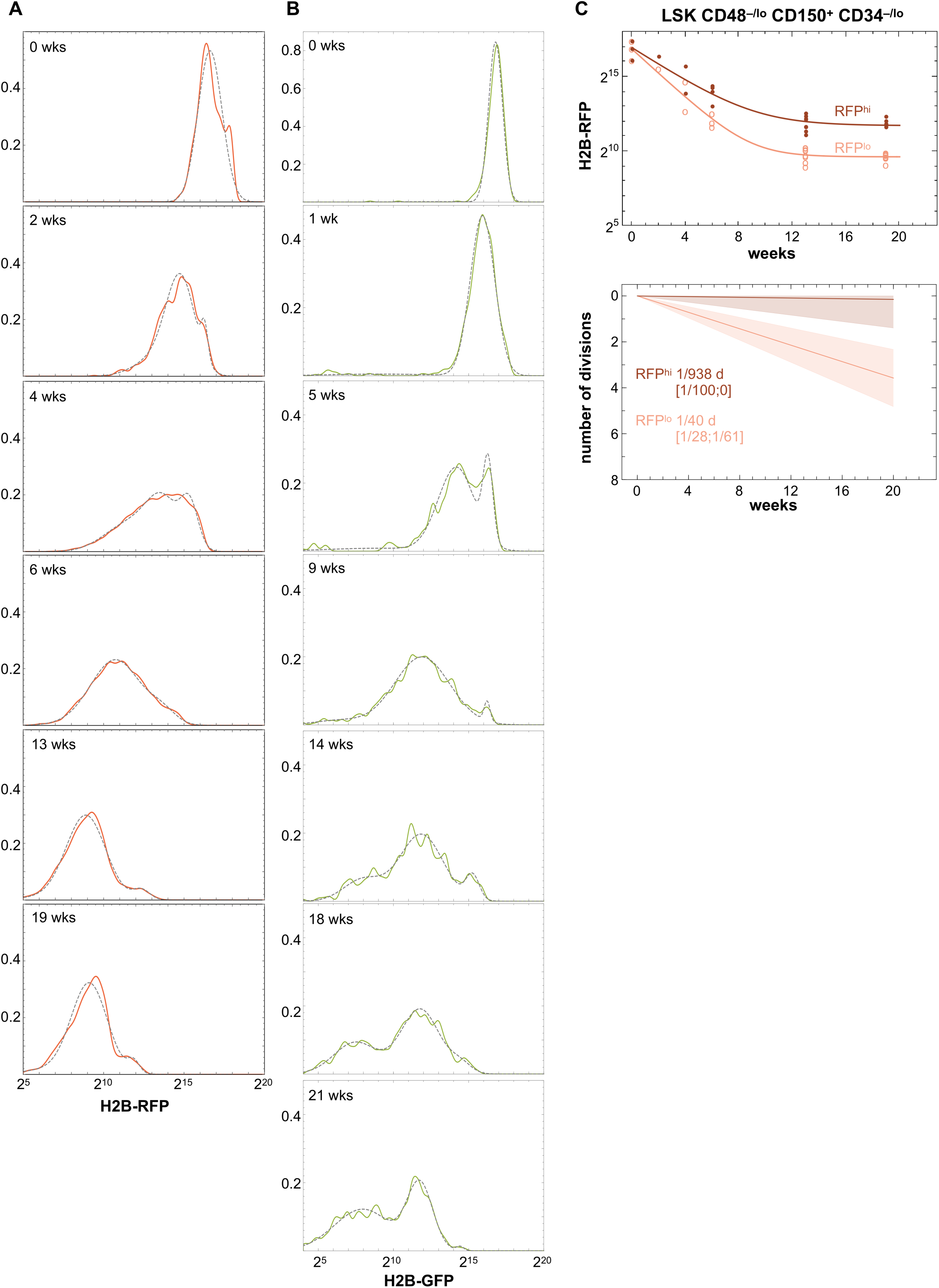
Mitotic activity of HSC populations inferred from H2B-FP label dilution data. **A-B** Representative examples for mixtures of Gaussian distributions, which were fitted to H2B-RFP (A, data from Morcos et al. 2017) or H2B-GFP (B, data from experiment shown in Figure 2A ‘GFP Morcos’) pulse-chase data of HSCs (continuous line: recorded data; dotted grey: fitted mixture distributions). Two Gaussians (RFP^lo^ and RFP^hi^) were fitted to H2B-RFP retention data (0-19 wks of chase), while for H2B-GFP data (0-21 wks of chase) two peaks (GFP^lo^ and GFP^hi^) were initially observed, but starting from 14 wks a third, intermediate peak uniformly appeared (GFP^mid^). **C** H2B-RFP dilution time course of fitted RFP^lo^ (light red, open circles) and RFP^hi^ (dark red, filled circles) Gaussian distributions from chased CD34^−/lo^ HSCs (LSK CD48^−/lo^CD150^+^CD34^−/lo^, upper panel, individual mice from Morcos et al. 2017) is shown. Division rates (lower panel) and respective 95 % confidence intervals (tinted area and square brackets) of RFP^lo^ and RFP^hi^ CD34^−/lo^ HSCs were calculated.

**Figure S3.**
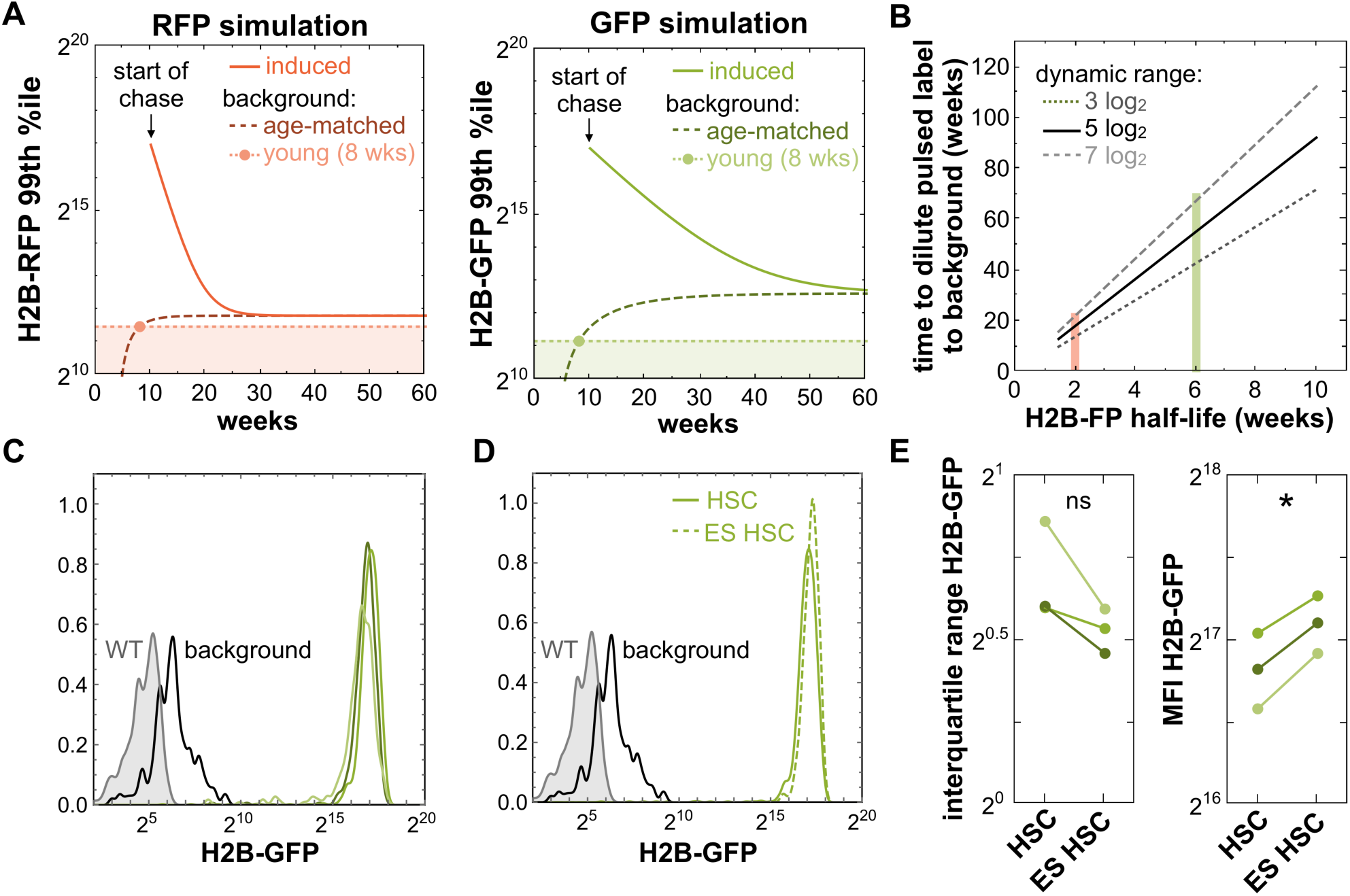
Dynamics of H2B-FP dilution experiments. **A** Simulation depicts H2B-FP pulse-chase experiments employing either H2B-RFP (left panel) or H2B-GFP (right panel) mouse models. Label decay in HSCs of pulsed and chased mice (continuous line) was calculated using our mathematical model. The age-dependent accumulation of background fluorescence in repressed control mice (dashed line) was additionally simulated. For the special case of a non-age matched background control (young, light dotted line), we arbitrary assumed 8 weeks of a age, which equals to ~4 weeks after HSCs have entered a predominantly quiescent state. Non-aged matched background controls result in under-estimation of the gating threshold for identification of H2B-FP^+^ HSCs and a seemingly label-retaining HSC population. **B** Simulation reveals the maximum chase interval until pulsed label fluorescence completely merges with accumulating background fluorescence. Merging of the populations was defined as the point when background fluorescence intensity equals 95% of the fluorescence intensity of the chased population. This simulation was carried out for 3 different arbitrary dynamic ranges (span between maximum pulse labeling and background fluorescence) and variable protein half-lives of the H2B-FP (half-life estimates for H2B-RFP and H2B-GFP are depicted in red and green, respectively). **C-E** *R26*^rtTA/rtTA^/*Col1A1*^H2B-GFP/H2B-GFP^ mice (n=3, shown in green) were DOX-induced (2g DOX/kg chow ad libitum) for three weeks. Un-induced *R26*^rtTA/rtTA^/*Col1A1*^H2B-GFP/H2B-GFP^ (black line) and B6 (WT, tinted grey area) mice served as background and auto-fluorescence controls, respectively. **C** Homogeneity of H2B-GFP labeling in HSCs (LSK CD48^−/lo^CD150^+^) in three pulsed individuals is depicted. **D** H2B-GFP distributions of LSK CD48^−/lo^CD150^+^ (HSC, continuous green line) and CD201^hi^ LS^hi^K CD48^−/lo^CD150^+^ (ES HSC, dashed green line) populations from a representative DOX-pulsed *R26*^rtTA/rtTA^/*Col1A1*^H2B-GFP/H2B-GFP^ mouse were overlaid and exemplify similar labeling homogeneity of HSCs and the primitive ES HSC subpopulation. **E** Interquartile range (left plot) and median (right plot) of H2B-GFP fluorescence intensity of the total HSC population were compared to the respective ES HSC subpopulation of the same DOX-induced individual (connecting lines). The interquartile ranges (labeling span of average events) of HSCs and ES HSCs revealed similar labeling homogeneity, while ES HSCs exhibited slightly higher labeling than HSCs.

**Table S1.**
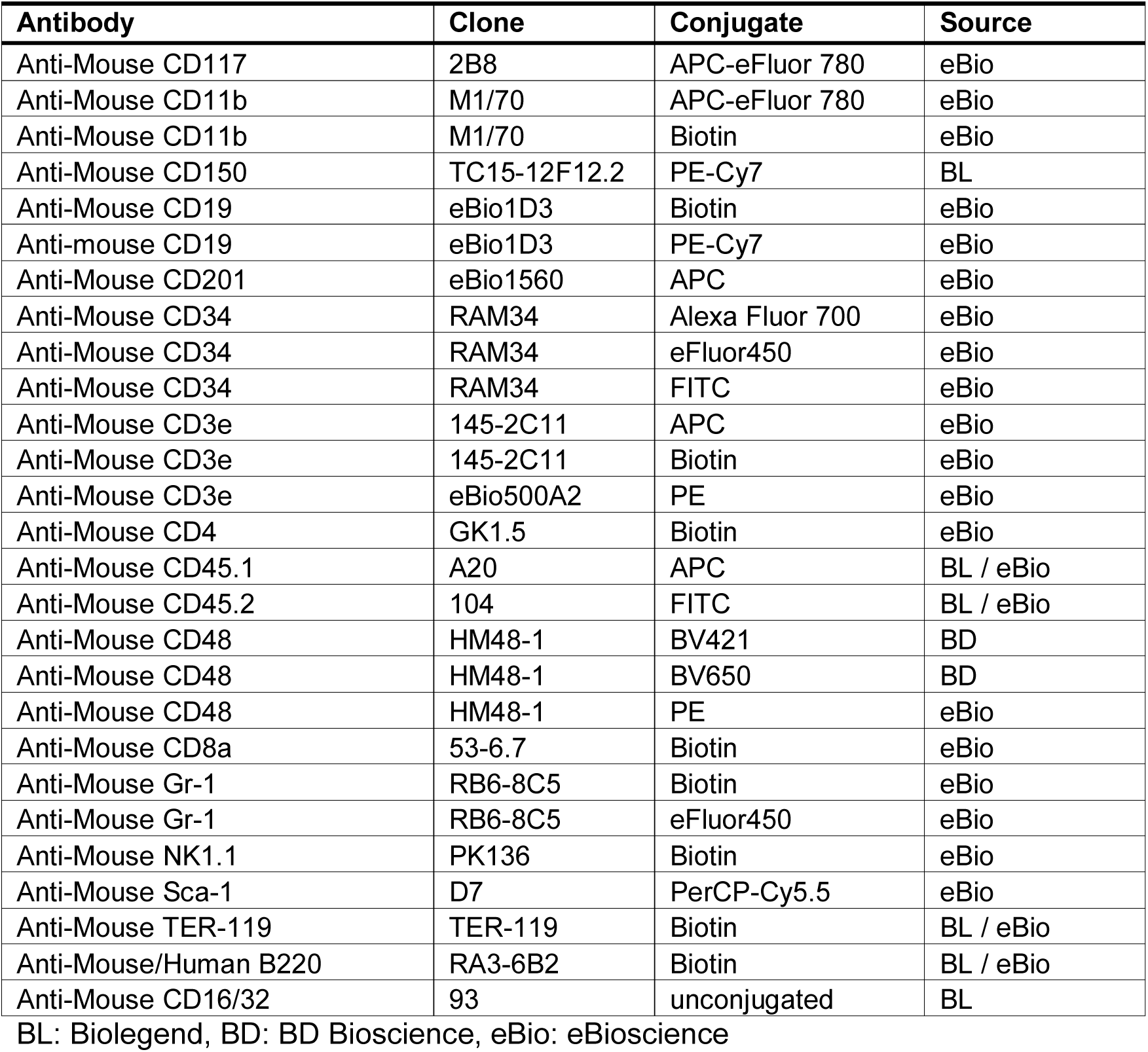
Antibody conjugates used for flow cytometry.

| Antibody | Clone | Conjugate | Source |
| --- | --- | --- | --- |
| Anti-Mouse CD117 | 2B8 | APC-eFluor 780 | eBio |
| Anti-Mouse CD11b | M1/70 | APC-eFluor 780 | eBio |
| Anti-Mouse CD11b | M1/70 | Biotin | eBio |
| Anti-Mouse CD150 | TC15-12F12.2 | PE-Cy7 | BL |
| Anti-Mouse CD19 | eBio1D3 | Biotin | eBio |
| Anti-mouse CD19 | eBio1D3 | PE-Cy7 | eBio |
| Anti-Mouse CD201 | eBio1560 | APC | eBio |
| Anti-Mouse CD34 | RAM34 | Alexa Fluor 700 | eBio |
| Anti-Mouse CD34 | RAM34 | eFluor450 | eBio |
| Anti-Mouse CD34 | RAM34 | FITC | eBio |
| Anti-Mouse CD3e | 145-2C11 | APC | eBio |
| Anti-Mouse CD3e | 145-2C11 | Biotin | eBio |
| Anti-Mouse CD3e | eBio500A2 | PE | eBio |
| Anti-Mouse CD4 | GK1.5 | Biotin | eBio |
| Anti-Mouse CD45.1 | A20 | APC | BL / eBio |
| Anti-Mouse CD45.2 | 104 | FITC | BL / eBio |
| Anti-Mouse CD48 | HM48-1 | BV421 | BD |
| Anti-Mouse CD48 | HM48-1 | BV650 | BD |
| Anti-Mouse CD48 | HM48-1 | PE | eBio |
| Anti-Mouse CD8a | 53-6.7 | Biotin | eBio |
| Anti-Mouse Gr-1 | RB6-8C5 | Biotin | eBio |
| Anti-Mouse Gr-1 | RB6-8C5 | eFluor450 | eBio |
| Anti-Mouse NK1.1 | PK136 | Biotin | eBio |
| Anti-Mouse Sca-1 | D7 | PerCP-Cy5.5 | eBio |
| Anti-Mouse TER-119 | TER-119 | Biotin | BL / eBio |
| Anti-Mouse/Human B220 | RA3-6B2 | Biotin | BL / eBio |
| Anti-Mouse CD16/32 | 93 | unconjugated | BL |

BL: Biolegend, BD: BD Bioscience, eBio: eBioscience

## References

Basak, O., M. van de Born, J. Korving, J. Beumer, S. van der Elst, J.H. van Es, and H. Clevers. 2014. Mapping early fate determination in Lgr5+ crypt stem cells using a novel Ki67-RFP allele. The EMBO Journal 33:2057–2068.

Beard, C., K. Hochedlinger, K. Plath, A. Wutz, and R. Jaenisch. 2006. Efficient method to generate single-copy transgenic mice by site-specific integration in embryonic stem cells. genesis 44:23–28.

Beerman, I., D. Bhattacharya, S. Zandi, M. Sigvardsson, I.L. Weissman, D. Bryder, and D.J. Rossi. 2010. Functionally distinct hematopoietic stem cells modulate hematopoietic lineage potential during aging by a mechanism of clonal expansion. Proceedings of the National Academy of Sciences 107:5465–5470.

Bernitz, J.M., H.S. Kim, B. MacArthur, H. Sieburg, and K. Moore. 2016. Hematopoietic Stem Cells Count and Remember Self-Renewal Divisions. Cell 167:1296–1309.e1210.

Bowie, M.B., K.D. McKnight, D.G. Kent, L. McCaffrey, P.A. Hoodless, and C.J. Eaves. 2006. Hematopoietic stem cells proliferate until after birth and show a reversible phase-specific engraftment defect. J Clin Invest 116:2808–2816.

Busch, K., K. Klapproth, M. Barile, M. Flossdorf, T. Holland-Letz, S.M. Schlenner, M. Reth, T. Höfer, and H.-R. Rodewald. 2015. Fundamental properties of unperturbed haematopoiesis from stem cells in vivo. Nature 518:542–546.

Challen, G.A., and M.A. Goodell. 2008. Promiscuous Expression of H2B-GFP Transgene in Hematopoietic Stem Cells. PLOS ONE 3:e2357.

Eaves, C.J. 2015. Hematopoietic stem cells: concepts, definitions, and the new reality. Blood 125:2605–2613.

Egli, D., J. Rosains, G. Birkhoff, and K. Eggan. 2007. Developmental reprogramming after chromosome transfer into mitotic mouse zygotes. Nature 447:679–685.

Foudi, A., K. Hochedlinger, D. Van Buren, J.W. Schindler, R. Jaenisch, V. Carey, and H. Hock. 2008. Analysis of histone 2B-GFP retention reveals slowly cycling hematopoietic stem cells. Nat. Biotechnol. 27:84–90.

Glauche, I., K. Moore, L. Thielecke, K. Horn, M. Loeffler, and I. Roeder. 2009. Stem cell proliferation and quiescence - two sides of the same coin. PLoS computational biology 5:e1000447.

Gossen, M., and H. Bujard. 2002. Studying Gene Function in Eukaryotes by Conditional Gene Inactivation. Annual Review of Genetics 36:153–173.

Grinenko, T., K. Arndt, M. Portz, N. Mende, M. Günther, K.N. Cosgun, D. Alexopoulou, N. Lakshmanaperumal, I. Henry, A. Dahl, and C. Waskow. 2014. Clonal expansion capacity defines two consecutive developmental stages of long-term hematopoietic stem cells. The Journal of Experimental Medicine 211:209–215.

Höfer, T., K. Busch, K. Klapproth, and H.-R. Rodewald. 2016. Fate Mapping and Quantitation of Hematopoiesis In Vivo. Annual Review of Immunology 34:449–478.

Huang, D.W., B.T. Sherman, and R.A. Lempicki. 2008. Systematic and integrative analysis of large gene lists using DAVID bioinformatics resources. Nature protocols 4:44.

Jamai, A., R.M. Imoberdorf, and M. Strubin. 2007. Continuous Histone H2B and Transcription-Dependent Histone H3 Exchange in Yeast Cells outside of Replication. Molecular Cell 25:345–355.

Kent, D.G., M.R. Copley, C. Benz, S. Wöhrer, B.J. Dykstra, E. Ma, J. Cheyne, Y. Zhao, M.B. Bowie, Y. Zhao, M. Gasparetto, A. Delaney, C. Smith, M. Marra, and C.J. Eaves. 2009. Prospective isolation and molecular characterization of hematopoietic stem cells with durable self-renewal potential. Blood 113:6342–6350.

Kiel, M.J., S. He, R. Ashkenazi, S.N. Gentry, M. Teta, J.A. Kushner, T.L. Jackson, and S.J. Morrison. 2007. Haematopoietic stem cells do not asymmetrically segregate chromosomes or retain BrdU. Nature 449:238–242.

Kiel, M.J., Ö.H. Yilmaz, T. Iwashita, O.H. Yilmaz, C. Terhorst, and S.J. Morrison. 2005. SLAM Family Receptors Distinguish Hematopoietic Stem and Progenitor Cells and Reveal Endothelial Niches for Stem Cells. Cell 121:1109–1121.

Kimura, H., and P.R. Cook. 2001. Kinetics of Core Histones in Living Human Cells. Little Exchange of H3 and H4 and Some Rapid Exchange of H2b 153:1341–1354.

Li, Q., N. Bohin, T. Wen, V. Ng, J. Magee, S.-C. Chen, K. Shannon, and S.J. Morrison. 2013. Oncogenic Nras has bimodal effects on stem cells that sustainably increase competitiveness. Nature 504:143.

McCarthy, D.J., K.R. Campbell, A.T.L. Lun, and Q.F. Wills. 2017. Scater: pre-processing, quality control, normalization and visualization of single-cell RNA-seq data in R. Bioinformatics (Oxford, England) 33:1179–1186.

Morcos, M.N.F., K.B. Schoedel, A. Hoppe, R. Behrendt, O. Basak, H.C. Clevers, A. Roers, and A. Gerbaulet. 2017. SCA-1 Expression Level Identifies Quiescent Hematopoietic Stem and Progenitor Cells. Stem Cell Reports 8:1472–1478.

Nacu, S., and T.D. Wu. 2010. Fast and SNP-tolerant detection of complex variants and splicing in short reads. Bioinformatics 26:873–881.

Nakada, D., H. Oguro, B.P. Levi, N. Ryan, A. Kitano, Y. Saitoh, M. Takeichi, G.R. Wendt, and S.J. Morrison. 2014. Oestrogen increases haematopoietic stem-cell self-renewal in females and during pregnancy. Nature 505:555.

Oguro, H., L. Ding, and Sean J. Morrison. 2013. SLAM Family Markers Resolve Functionally Distinct Subpopulations of Hematopoietic Stem Cells and Multipotent Progenitors. Cell Stem Cell 13:102–116.

Osawa, M., K. Hanada, H. Hamada, and H. Nakauchi. 1996. Long-term lymphohematopoietic reconstitution by a single CD34-low/negative hematopoietic stem cell. Science 273:242–245.

Passegué, E., A.J. Wagers, S. Giuriato, W.C. Anderson, and I.L. Weissman. 2005. Global analysis of proliferation and cell cycle gene expression in the regulation of hematopoietic stem and progenitor cell fates. The Journal of Experimental Medicine 202:1599–1611.

Qiu, J., D. Papatsenko, X. Niu, C. Schaniel, and K. Moore. 2014. Divisional History and Hematopoietic Stem Cell Function during Homeostasis. Stem Cell Reports 2:473–490.

Radomska, H.S., D.A. Gonzalez, Y. Okuno, H. Iwasaki, A. Nagy, K. Akashi, D.G. Tenen, and C.S. Huettner. 2002. Transgenic targeting with regulatory elements of the human *CD34* gene. Blood 100:4410–4419.

Sato, T., J.H. Laver, and M. Ogawa. 1999. Reversible Expression of CD34 by Murine Hematopoietic Stem Cells. Blood 94:2548–2554.

Säwén, P., S. Lang, P. Mandal, D. Rossi, S. Soneji, and D. Bryder. 2016. Mitotic History Reveals Distinct Stem Cell Populations and Their Contributions to Hematopoiesis. Cell Reports 14:2809–2818.

Schoedel, K., M. Morcos, T. Zerjatke, I. Roeder, T. Grinenko, D. Voehringer, J. Göthert, C. Waskow, A. Roers, and A. Gerbaulet. 2016. The bulk of the hematopoietic stem cell population is dispensable for murine steady-state and stress hematopoiesis. Blood 128:2285–2296.

Sheikh, B.N., Y. Yang, J. Schreuder, S.K. Nilsson, R. Bilardi, S. Carotta, H.M. McRae, D. Metcalf, A.K. Voss, and T. Thomas. 2016. MOZ (KAT6A) is essential for the maintenance of classically defined adult hematopoietic stem cells. Blood 128:2307–2318.

Shin, J.Y., W. Hu, M. Naramura, and C.Y. Park. 2014. High c-Kit expression identifies hematopoietic stem cells with impaired self-renewal and megakaryocytic bias. The Journal of Experimental Medicine 211:217–231.

Smyth, G.K., W. Shi, and Y. Liao. 2014. featureCounts: an efficient general purpose program for assigning sequence reads to genomic features. Bioinformatics 30:923–930.

Sun, J., A. Ramos, B. Chapman, J.B. Johnnidis, L. Le, Y.-J. Ho, A. Klein, O. Hofmann, and F.D. Camargo. 2014. Clonal dynamics of native haematopoiesis. Nature 514:322–327.

Takizawa, H., R.R. Regoes, C.S. Boddupalli, S. Bonhoeffer, and M.G. Manz. 2011. Dynamic variation in cycling of hematopoietic stem cells in steady state and inflammation. J Exp Med 208:273–284.

Toyama, Brandon H., Jeffrey N. Savas, Sung K. Park, Michael S. Harris, Nicholas T. Ingolia, John R. Yates, and Martin W. Hetzer. 2013. Identification of Long-Lived Proteins Reveals Exceptional Stability of Essential Cellular Structures. Cell 154:971–982.

Tumbar, T., G. Guasch, V. Greco, C. Blanpain, W.E. Lowry, M. Rendl, and E. Fuchs. 2004. Defining the Epithelial Stem Cell Niche in Skin. Science 303:359–363.

van der Wath, R.C., A. Wilson, E. Laurenti, A. Trumpp, and P. Liò. 2009. Estimating Dormant and Active Hematopoietic Stem Cell Kinetics through Extensive Modeling of Bromodeoxyuridine Label-Retaining Cell Dynamics. PLOS ONE 4:e6972.

Venkatesh, S., and J.L. Workman. 2015. Histone exchange, chromatin structure and the regulation of transcription. Nature Reviews Molecular Cell Biology 16:178.

Waghmare, S.K., R. Bansal, J. Lee, Y.V. Zhang, D.J. McDermitt, and T. Tumbar. 2008. Quantitative proliferation dynamics and random chromosome segregation of hair follicle stem cells. The EMBO Journal 27:1309–1320.

Wilson, A., E. Laurenti, G. Oser, R.C. van der Wath, W. Blanco-Bose, M. Jaworski, S. Offner, C.F. Dunant, L. Eshkind, E. Bockamp, P. Lió, H.R. MacDonald, and A. Trumpp. 2008. Hematopoietic Stem Cells Reversibly Switch from Dormancy to Self-Renewal during Homeostasis and Repair. Cell 135:1118–1129.

Wilson, N., D. Kent, F. Buettner, M. Shehata, I. Macaulay, F. Calero-Nieto, M. Sanchez Castillo, C. Oedekoven, E. Diamanti, R. Schulte, C. Ponting, T. Voet, C. Caldas, J. Stingl, A. Green, F. Theis, and B. Göttgens. 2015. Combined Single-Cell Functional and Gene Expression Analysis Resolves Heterogeneity within Stem Cell Populations. Cell Stem Cell 16:712–724.

Zerbino, D.R., P. Achuthan, W. Akanni, M.R. Amode, D. Barrell, J. Bhai, K. Billis, C. Cummins, A. Gall, C.G. Girón, L. Gil, L. Gordon, L. Haggerty, E. Haskell, T. Hourlier, O.G. Izuogu, S.H. Janacek, T. Juettemann, J.K. To, M.R. Laird, I. Lavidas, Z. Liu, J.E. Loveland, T. Maurel, W. McLaren, B. Moore, J. Mudge, D.N. Murphy, V. Newman, M. Nuhn, D. Ogeh, C.K. Ong, A. Parker, M. Patricio, H.S. Riat, H. Schuilenburg, D. Sheppard, H. Sparrow, K. Taylor, A. Thormann, A. Vullo, B. Walts, A. Zadissa, A. Frankish, S.E. Hunt, M. Kostadima, N. Langridge, F.J. Martin, M. Muffato, E. Perry, M. Ruffier, D.M. Staines, S.J. Trevanion, B.L. Aken, F. Cunningham, A. Yates, and P. Flicek. 2017. Ensembl 2018. Nucleic Acids Research 46:D754–D761.

